# A Locally Activatable Sensor for Robust Quantification of Organellar Glutathione

**DOI:** 10.1101/2022.04.01.486692

**Authors:** Sarah Hübner, Gianluca Quargnali, Sebastian Thallmair, Pablo Rivera-Fuentes

## Abstract

Glutathione (GSH) is the main determinant of intracellular redox potential and participates in multiple cellular signaling pathways. Achieving a detailed understanding of intracellular GSH trafficking and regulation depends on the development of tools to map GSH compartmentalization and intra-organelle fluctuations. Herein, we present a new GSH sensing platform, TRaQ-G, for live-cell imaging. This small-molecule/protein hybrid sensor possesses a unique reactivity turn-on mechanism that ensures that the small molecule is only sensitive to GSH in the desired location. Furthermore, TRaQ-G can be fused to a fluorescent protein of choice to give a ratiometric response. Using TRaQ-G-mGold, we demonstrated that the nuclear and cytosolic GSH pools are independently regulated during cell proliferation. We also used this sensor, in combination with roGFP, to quantify redox potential and GSH concentration simultaneously in the endoplasmic reticulum. Finally, by exchanging the fluorescent protein, we created a near-infrared, targetable and quantitative GSH sensor.

## 1 Introduction

Glutathione (GSH) is a small peptide (L-γ-glutamyl-L-cysteinyl glycine) that can react as a nucleophile or a reducing agent. Upon oxidation, it forms a disulfide-bonded dimer (GSSG). Owing to its high concentration in the cell (∼10^−3^ M)^1^, the GSH/GSSG pair acts as the main intracellular redox buffer, plays a key role in detoxification of reactive oxygen species (ROS), and participates in redox signaling^2^. Furthermore, the concentration of GSH and the GSH/GSSG ratio vary between intracellular organelles and this compartmentalization is actively regulated in the cell^3^. Consequently, disruption of intracellular GSH homeostasis is linked to many pathologies, including cancer^4^, diabetes^5^, and neurodegenerative disorders^6^. A detailed understanding of the mechanisms of intracellular redox homeostasis and their connection to disease depends on our ability to quantify, among other parameters, changes in GSH concentration in specific organelles. Therefore, there is a need for robust tools that can quantify this critical redox modulator with subcellular specificity.

The GSH/GSSG ratio can be measured in specific organelles using redox-sensitive green fluorescent proteins (roGFPs), particularly those fused to the GSH-specific Grx1 protein^7^. Despite their usefulness, these proteins operate at short wavelengths (<450 nm), display modest brightness, and have limited dynamic ranges^8^. Small-molecule sensors have been a popular choice to measure total GSH concentrations. Most of these probes are π-conjugated electrophiles that change their fluorescence upon reaction with GSH. Notable examples include coumarin^9–12^ and silicon rhodamine (SiR)^13^ dyes. The latter are particularly useful because of their red-shifted excitation, selectivity, reversibility, and fast kinetics. Targeting small molecules to specific subcellular compartments, however, is challenging. In the case of GSH sensing, the problem is even more difficult because in order to reach its intended organelle, the small molecule must cross the cytosol, where the concentration of GSH is in the 10^−3^ M range. Thus, sensors could react with GSH in the cytosol, and it is difficult to assess whether small molecules actually sense the real concentrations of GSH in their target organelle or shuttle GSH from the cytosol. In fact, a similar question could be asked of any other targeted small-molecule fluorescent sensor. With this limitation in mind, we set out to develop a GSH sensor that is locally activated in the compartment of interest, thus providing a reliable measure of intra-organelle GSH concentrations.

We based our design on the reactivity of SiR dyes, which in their zwitterionic form can serve as electrophiles that react with GSH (Figure 1a)^13^. We hypothesized that a SiR dye in a spirocyclic form would be non-reactive against GSH (Figure 1b), but it could isomerize to its zwitterionic form upon binding to the self-labeling protein HaloTag (HT)^14–16^, switching both its fluorescence and reactivity against GSH from an “off” to an “on” state. Moreover, this hybrid sensor design combines the advantages of genetically encoded sensors, such as precise targetability, with the tunability, brightness, and photostability of small molecules. Finally, by fusing a fluorescent protein (FP) to HT, we could have an internal standard for ratiometric measurements. We call this design **T**argetable **Ra**tiometric **Q**uantitative **G**SH (TRaQ-G) probes (Figure 1c).

**Figure 1.**
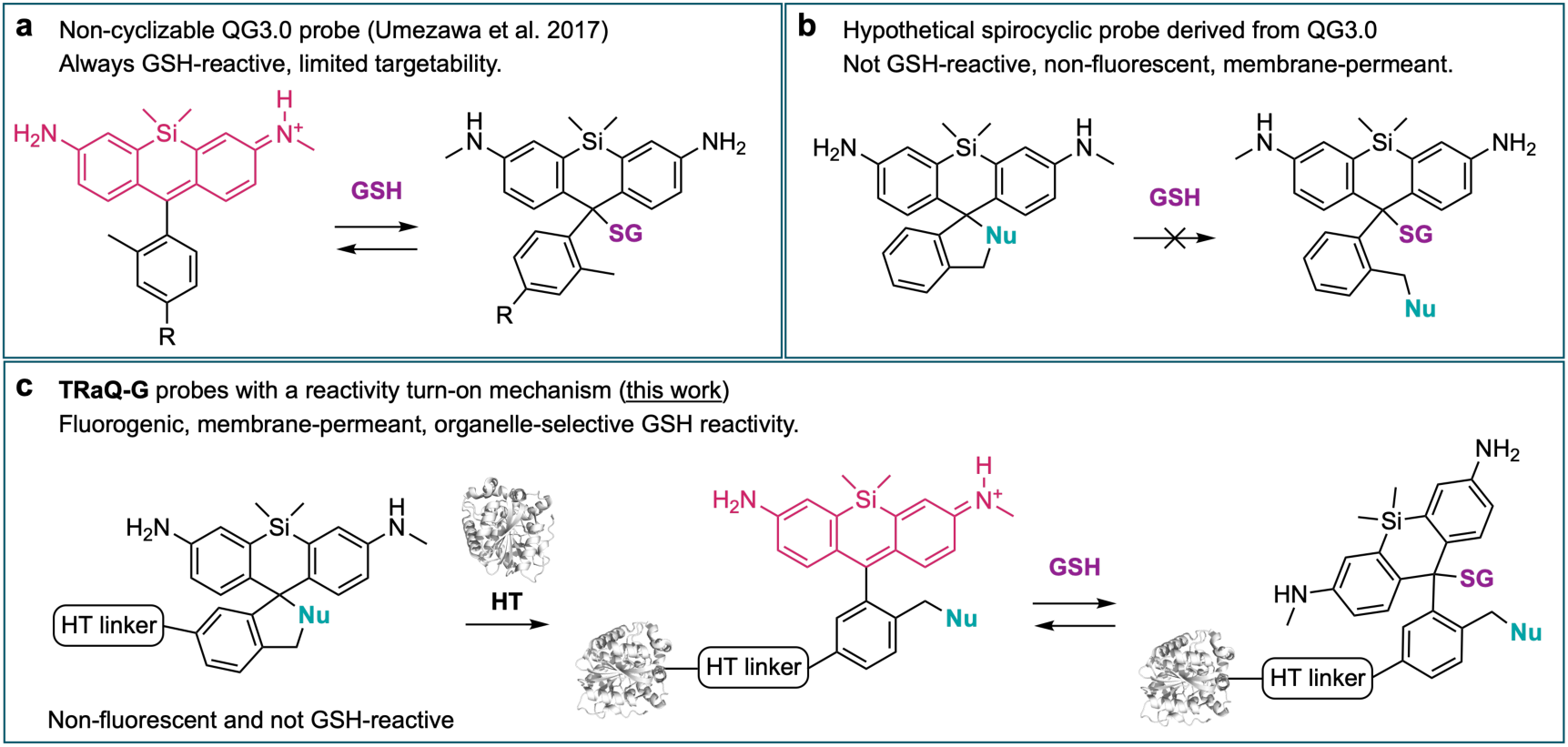
Previously reported SiR-based GSH sensors and TRaQ-G sensing concept. R = short linker and secondary small-molecule fluorophore for ratiometric imaging^13^. HT linker = chloroalkane substrate.

## 2 Results and Discussion

### Structure-reactivity analysis of TRaQ-G probes

Our first goal was to demonstrate that a known SiR-based GSH sensor, QG3.0^13^, retained its reactivity towards GSH when bound to HT. We therefore substituted the reference fluorophore of QG3.0 with a chloroalkane linker, a substrate that reacts with HT (Figure 2a). The SiR unit was synthesized according to the reported procedure^13^ and the linker was attached *via* amide coupling (Scheme S1) to give the Me-TRaQ-G ligand. Using this molecule, we prepared and purified the HT adduct (see Methods section), Me-TRaQ-G. The reactivity of Me-TRaQ-G against GSH was nearly identical to that of QG3.0 (Figure 2c and Extended Data Figure 1a-c)^13^ suggesting that one face of the xanthene π-system of Me-TRaQ-G must be exposed to the solvent, thus allowing for GSH nucleophilic attack. This hypothesis was confirmed by X-ray diffraction studies of the adduct, which revealed that the methyl group of the Me-TRaQ-G ligand fits into a shallow hydrophobic pocket created by residues F152, V167 and T172 of HT, exposing the opposite face of the xanthene core to the solvent (Figure 2b). Furthermore, molecular dynamics (MD) simulations based on the crystal structure indicate that this orientation of the ligand is preferred over the conformation in which the methyl group is exposed to the solvent (Extended Data Figure 1d-e).

**Figure 2.**
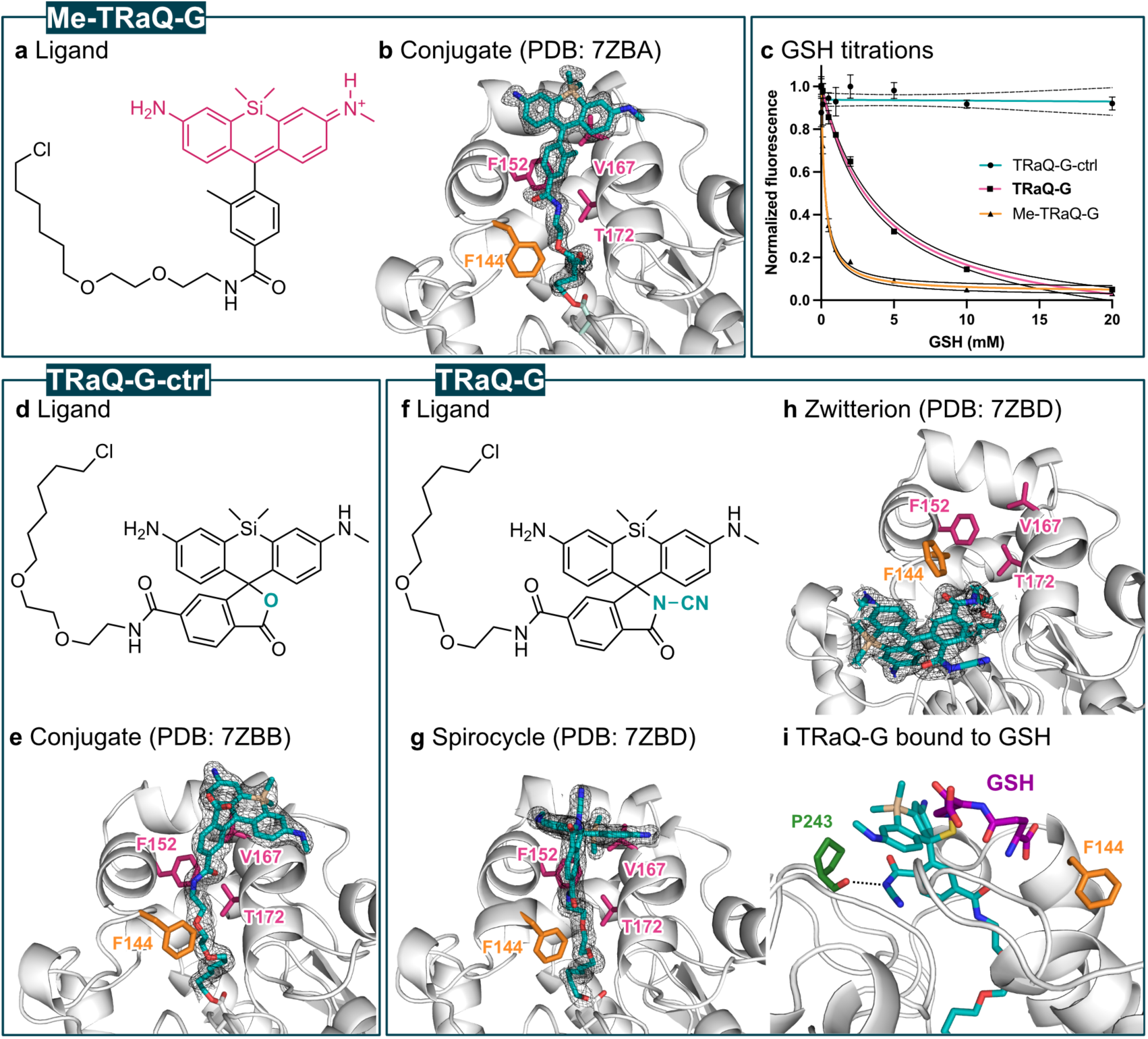
Structures and reactivity of TRaQ-G probes. a) Chemical structure of the small-molecule ligand for Me-TRaQ-G. b) X-ray crystal structure of Me-TraQ-G (1.23 Å). c) GSH titrations of the three TRaQ-G probes. The concentration of the TRaQ-G probe was kept at 5–15 µM. d) Chemical structure of the small-molecule ligand for TRaQ-G-ctrl. e) X-ray crystal structure of TraQ-G-ctrl (1.95 Å). f) Chemical structure of the small-molecule ligand for TRaQ-G. g) X-ray crystal structure of TraQ-G (1.69 Å) displaying the SiR ligand in the spirocyclic form, and h) the zwitterionic form. i) Snapshot at 500 ns of the MD simulation of TRaQ-G with GSH bound. Residues forming the hydrophobic pocket are displayed in pink, the Phe residue that moves between the open and closed conformation is displayed in orange and the residue forming a hydrogen bond with the ligand is displayed in green. The 2Fo-Fc electron density map is displayed around the ligand with σ=1.

Next, we tested whether SiR derivatives that undergo spirocyclization could become reactive against GSH upon binding to HT. For this purpose, we prepared derivatives in which either a carboxylic acid (Figure 2d)^14^ or a cyanamide (Figure 2f)^16^ nucleophile could form the spirocycle. The syntheses of these probes were adapted from published procedures (Scheme S2 and Scheme S3)^14,16^.

Both ligands were obtained in their spirocyclic form and reacted only very slowly (TRaQ-G) or not at all (TRaQ-G-ctrl) with GSH, as judged by mass spectrometry (Figure S1). Upon reaction with HT, both ligands produced intensely blue solutions, indicating that the zwitterionic form of the SiR core is formed. GSH titration experiments revealed that, whereas the HT-bound carboxylic acid derivative was completely insensitive to GSH, the cyanamide ligand bound to HT responded to GSH in the physiologically relevant range of concentrations (∼1– 20 mM, Figure 2c). We further confirmed that this probe is susceptible only to the absolute GSH concentration, not to the GSH/GSSG ratio (Figure S2a), and that GSH binding is reversible (Figure S2b). Furthermore, no other common intracellular nucleophiles reacted with the probe at physiologically relevant concentrations (Figure S2c). We envisioned that the carboxylic acid derivative, which is insensitive to GSH, could be used as a control probe in quantitative studies. Therefore, we refer to the HT-bound carboxylic acid derivative as TRaQ-G-ctrl, whereas the cyanamide derivative is hereafter called TRaQ-G.

To understand the difference in reactivity between TRaQ-G-ctrl and TRaQ-G, we analyzed the structure of both conjugates by X-ray diffraction. In TRaQ-G-ctrl, the xanthene core of the SiR dye rests on top of the hydrophobic surface created by residues F152, V167 and T172 of HT, and the carboxylate group points outwards into the solvent (Figure 2e). MD simulations indicate that this conformation is very stable, arguably because of solvation of the carboxylate moiety and additional hydrogen bonding from the secondary amine of the ligand to residue E170 (Figure S3). This carboxylate, and its solvation sphere, efficiently block GSH attack, as observed in GSH titrations (Figure 2c). In the case of TRaQ-G, we found two distinct monomers in the unit cell, one in which the ligand is in the spirocyclic form (Figure 2g) and the other in which the xanthene is in the zwitterionic form (Figure 2h). Whereas the spirocyclic form binds similarly to Me-TRaQ-G and TRaQ-G-ctrl, the zwitterionic form occupies an alternative pocket. MD simulations using the spirocyclic form as starting point suggest that the side chain of residue F144 must rotate to allow for the ligand to occupy the alternative pocket and thus switch to its zwitterionic form (Figure S4, Movie S1). Although mutation of this residue does not seem to increase the brightness of SiR-carboxylate dyes^17^, this residue could be important to create HT mutants that induce larger fluorogenicity in other SiR-based spirocyclic dyes. Using MD simulations starting from the zwitterionic structure, we found that the structure rapidly relaxes to a minimum in which the cyanamide forms a persistent hydrogen bond with the backbone carbonyl of residue P243 (Extended Data Figure 2). This interaction exposes the opposite face of the SiR dye to the solvent, allowing for GSH attack (Figure 2i), explaining the reactivity observed in titration experiments (Figure 2c).

### TRaQ-G is a robust, targetable and quantitative GSH sensor

Although Me-TRaQ-G is sensitive towards GSH, we envisioned that it would perform poorly in live cells given its constant fluorescence and reactivity towards GSH. This suspicion was confirmed by imaging cells expressing HT in the nucleus, which revealed unacceptable levels of off-target signals (Figure S5). Instead, we focused on TRaQ-G, the ligand of which undergoes fluorescence and reactivity turn-on upon binding to HT. Once bound to HT, the strong fluorescence of the TRaQ-G ligand decreases upon reaction with GSH. This turn-off behavior can be converted into a ratiometric readout by the presence of an internal standard that is insensitive to GSH. For this purpose, we chose the monomeric, bright, photostable, and redox-insensitive protein mGold^18^ to create a fusion with HT. Furthermore, this construct could be targeted to specific organelles using established peptide localization sequences (Figure 3a). In order to relate the TRaQ-G-mGold fluorescence ratio to a concentration of GSH, we expressed and purified the fusion protein HT-mGold and treated it with the TRaQ-G ligand. Mass spectrometry revealed that formation of the adduct is quantitative in vitro (Extended Data Figure 3a). After purification, this adduct was used to generate a calibration curve, relating the ratio of mGold/TRaQ-G fluorescence intensities to GSH concentrations (Extended Data Figure 3b). This ratiometric sensor responded linearly in the range 1–20 mM GSH, confirming its suitability to quantify concentrations of GSH in the physiological range.

**Figure 3.**
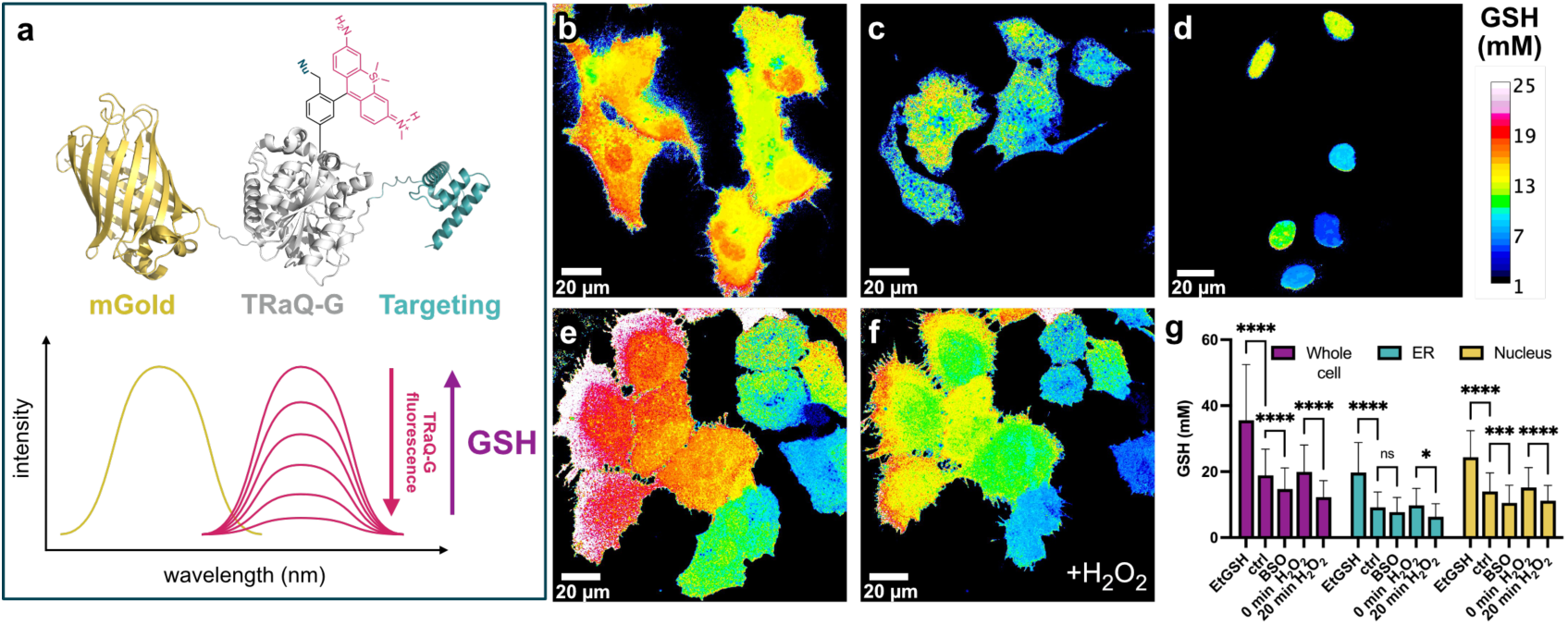
GSH quantification in subcellular locations using TRaQ-G-mGold. a) Schematic depiction of the GSH sensor TRaQ-G-mGold with a targeting peptide and mechanism of ratiometric sensing. Ratiometric imaging of GSH in whole cells (b), ER (c) and nuclei (d) using TRaQ-G-mGold. GSH quantification in cells before (e) and 20 min after addition of 1 mM H_2_O_2_ (f). g) Quantification of GSH concentration in whole cells, the ER or nuclei untreated (ctrl), after addition of 10 mM EtGSH (membrane-permeant GSH precursor), 1 mM BSO (inhibitor of GSH biosynthesis), or 1 mM H_2_O_2_ (oxidant). TRaQ-G ligand concentration = 100 nM in all cases. Images were generated by dividing the intensity of mGold over TRaQ-G and relating this ratio to the calibration curve obtained with purified TRaQ-G-mGold. For display purposes, calibrated images were despeckled using Fiji (ImageJ). Statistical significance was evaluated by two-way ANOVA (see Methods section for details) with *N* = 130, 132, 142, 111, 103, 118, 116, 131, 93, 90, 122, 149, 176, 143, 137 (from left to right) from three biological replicates. Mean and standard deviations are depicted and ns = P>0.05, * = P≤0.05, ** = P≤0.01, *** = P≤0.001, **** = P≤0.0001.

To test the applicability of TRaQ-G in live-cell imaging, we generated a set of plasmids encoding for HT-mGold in the whole cell, Calnexin-HT-mGold for endoplasmic reticulum (ER) targeting and H2B-HT-mGold for nuclear targeting (see Methods section). We transfected cells with these plasmids and treated them with 100 nM TRaQ-G ligand. With a short and simple incubation protocol, the sensor could be detected in the intended organelle with essentially no background signal (Figure 3b-d). The ligands can also be used without washing. To image the ER, which has a more oxidizing environment than the cytosol^19^, we also expressed a fusion of the new, redox-insensitive HT8^20^. Comparison of the redox-insensitive HT8 and the HT7 variant used for TRaQ-G indicated no measurable difference in binding efficiency or fluorogenicity (Figure S6a). We also measured the kinetics of labeling and found that intracellular labeling of HT is fast and reaches saturation in less than 30 min (Figure S6b). This experiment confirms that, under our imaging conditions, HT is stably labeled to its maximum extent. Furthermore, because the ligand does not have to be washed, newly expressed protein can continue to be labeled during long-term experiments.

Using TRaQ-G-mGold, we measured the equilibrium concentrations in the whole cell, the ER, and the nucleus. The concentration of GSH in the whole cell was approximately 18.9±9.4 mM. This figure is higher than previously reported values of ≤10 mM^9,21^, but existing probes have limited dynamic ranges, rarely exceeding 10 mM. In the ER, GSH concentrations were also higher, as observed before^19^, with values around 9.2±5.3 mM. Nuclei displayed GSH concentrations as high as 14.0±6.7 mM. We observed significant cell-to-cell variability in our GSH quantification experiments. These differences, in particular in the nuclear GSH pool, could be a consequence of cells being in different stages of the cell cycle^22^. This effect will be described in the next section.

Next, we tested whether mGold-TRaQ-G could be used to monitor changes in GSH concentration. For that purpose, we exposed cells equipped with the TRaQ-G-mGold sensor to reagents that would modify the intracellular GSH concentration (Figure 3e-g). All three subcellular regions displayed higher GSH concentrations after treatment with glutathione monoethyl ester (EtGSH), a cell-permeable precursor of GSH (Figure 3g). We hindered GSH biosynthesis by inhibiting γ-glutamylcysteine synthetase with buthionine sulfoximine (BSO), which led us to observe lower GSH concentrations in the whole cell and the nucleus, but not in the ER, at least during our observation times (3-4 h post-treatment). Finally, cells treated with 1 mM H_2_O_2_ (20 min) also displayed significant decrease in GSH concentrations across organelles (Figure 3e-g).

### Nuclear and cytosolic GSH pools are independently regulated during cell proliferation

The high cell-to-cell variability observed in nuclear GSH concentrations led us to hypothesize that these differences could be correlated with the phase of the cell cycle. Although the existence of an independent nuclear GSH pool has been debated in the literature^9,23^, there are indications that high GSH concentrations in the nucleus are required for cell proliferation and that the nuclear GSH pool varies during the cell cycle^22,23^. In particular, indirect evidence suggests that cells accumulate GSH in the nucleus in the S and G2 phases, prior to mitosis^22^. To test this hypothesis, we transfected HeLa cells with H2B-HT-mGold, synchronized them in the early S phase by double thymidine block^24^, incubated them with either TRaQ-G ligand or the GSH-insensitive TRaQ-G-ctrl ligand, and imaged them for 24 h. Ratiometric imaging with H2B-TRaQ-G-mGold indicated that GSH concentration in the nucleus increased slightly in the first 1-2 h after thymidine block release, which corresponds to early S phase (Figure 4a,c). After this time, GSH levels dropped steadily during S phase, staying more or less stable through G2 phase and mitosis (Figure4a,c). We employed probe H2B-TRaQ-G-ctrl-mGold, which is insensitive to GSH (Figure 2c), to demonstrate that these changes indeed reflect fluctuations in GSH concentration (Figure 4b,c) and are not a consequence of photobleaching, changes in expression levels of the sensor, or other labeling artifacts.

**Figure 4.**
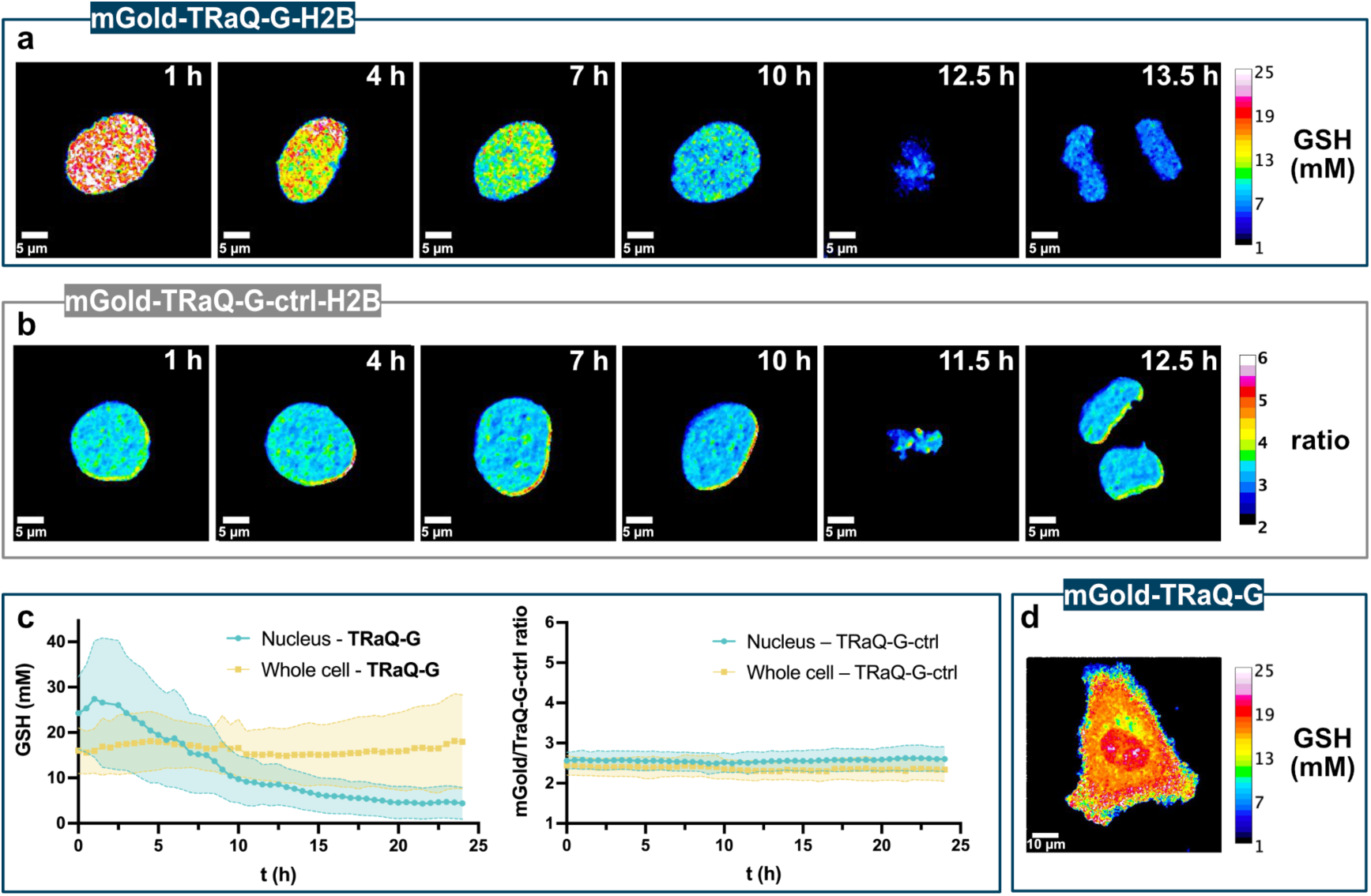
Regulation of nuclear GSH concentrations during cell proliferation. a) TRaQ-G-mGold ratiometric imaging of a single nucleus from S phase to mitosis, revealing a steady decrease in GSH concentration. b) Control probe TRaQ-G-ctrl-mGold confirms that the observed variations are a consequence of GSH fluctuations. c) Quantification of GSH concentrations or variation of ratio of control probe for 24 h after release of cells from thymidine block (early S-phase). Lines represent average and shaded areas standard deviation from *N* ≥ 19, 50, 42, 19 cells per timepoint from three biological replicates, for H2B-TRaQ-G-mGold, H2B-TRaQ-G-ctrl-mGold, TRaQ-G-mGold and TRaQ-G-ctrl-mGold, respectively. d) TRaQ-G-mGold ratiometric imaging of a single cell (t = 2 h) displaying different concentrations in the nucleus and the rest of the cell.

Next, we tested whether the high GSH concentration in S phase and subsequent decrease during G2 phase was observed only in the nucleus or in the whole cell. We transfected HeLa cells with HT-mGold and labeled them with either TRaQ-G or TRaQ-G-ctrl ligands. TRaQ-G imaging revealed that the GSH concentration in the whole cell fluctuates significantly less than in the nucleus (Figure 4c), and the concentration in the nucleus appears to be different to that in the rest of the cell (Figure 4d). It should be noted that GSH fluctuations for the whole cell were heterogenous between biological replicates, which may be a consequence of varying passage numbers or slightly different culture densities^7,22^, however, compartmentalization of nuclear GSH was observed in many cases. Finally, control experiments with TRaQ-G-ctrl in the whole cell indicated no noticeable variations in ratios between the nucleus or the rest of the cell throughout the experiment (Figure 4c).

These findings indicate that the GSH concentration in the nucleus is independently regulated from that elsewhere in the cell, is highest during S phase, and steadily decreases until mitosis. Furthermore, the existence of different GSH concentrations in the nucleus and the cytosol strongly implies that GSH does not diffuse freely across the nuclear envelope or through nuclear pores. Further studies, beyond the scope of this work, will be required to determine the molecular mechanisms of GSH regulation in the nucleus.

### Multicolor imaging with a roGFP and near-infrared GSH imaging

We envisioned that TRaQ-G-mGold could be multiplexed with roGFPs to measure the total GSH concentration and the GSH/GSSG ratio simultaneously. roGFPs are ratiometric sensors that undergo an excitation wavelength shift from 405 to about 445 nm upon reaction with GSSG^7,19^. Their emission wavelength is in both cases around 480 nm. Considering that mGold can be efficiently excited at 515 nm and its emission occurs around 570 nm, and TRaQ-G can be excited at 561 nm and emits at wavelengths beyond 600 nm, we planned the four-color experiment depicted in Figure 5a. Next, we co-transfect roGFP-iE-ER^19^, a roGFP variant that has been engineered for the ER, and Calnexin-TRaQ-G-mGold. Four-color imaging revealed that high GSH concentrations do not always correlate with a high GSH/GSSG ratio (Figure 5b,c). This observation suggests that even under conditions of high GSH accumulation, the concentration of GSSG also increases to maintain the redox potential. To test this hypothesis, we incubated cells with EtGSH, a membrane-permeant precursor of GSH. Using roGFP-iE-ER and Calnexin-TRaQ-G-mGold, we observed that the concentration of GSH increases significantly, but the redox potential becomes only marginally more reductive (Figure 5d). Although further experiments would be needed to understand this effect, these results demonstrate that TRaQ-G can be used in combination with roGFPs to study details of redox homeostasis in intracellular organelles.

Finally, the flexible design of TRaQ-G sensors allows for using a more red-shifted fluorescence protein. To create a targeted GSH sensor reaching into the near-infrared (NIR) region of the spectrum, we created a plasmid encoding for an H2B-HT-emiRFP703^25^ fusion protein. We transiently transfected HeLa cells with this construct, incubated them with 100 nM TRaQ-G and treated them with 1 mM H_2_O_2_. A drop in the emiRFP703/TRaQ-G ratio was observed corresponding to decreased GSH concentration. In particular, cells that had an initially high concentration (> 4 emiRFP703/TRaQ-G) showed a more pronounced change. A reason for this behavior could be that H_2_O_2_ does not oxidize GSH directly, but rather through the action of glutathione peroxidase^26^. This enzyme follows first-order kinetics with respect to GSH concentration and thus the initial reaction rate varies linearly with GSH molarity^27^. This consideration explains why under excess H_2_O_2_, cells with an initially higher GSH concentration display a faster GSH oxidation. However, additional studies would be necessary to validate these conclusions.

**Figure 5.**
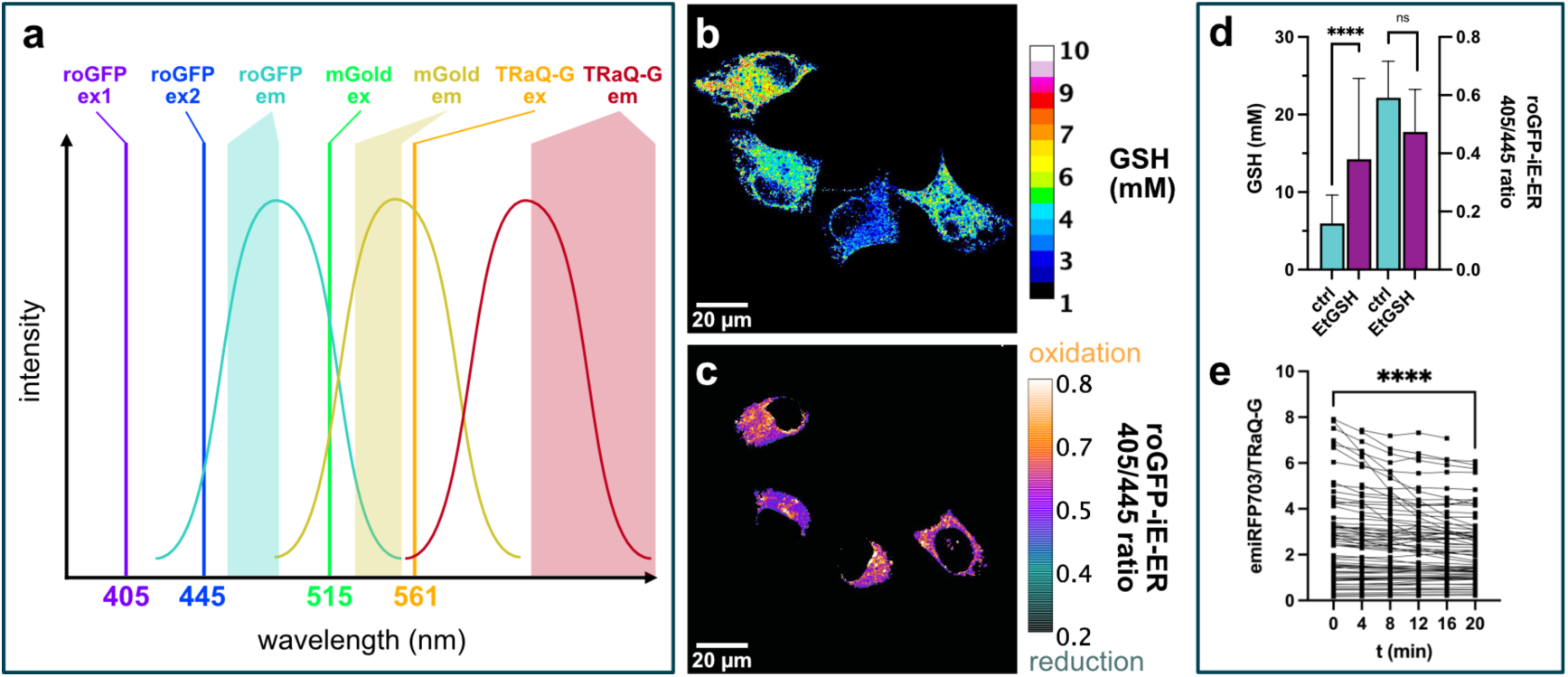
Multicolor and NIR imaging with TRaQ-G. a) Schematic representation of a four-color experiment multiplexing roGFP-iE-ER and Calnexin-TRaQ-G-mGold. Colored lines represent the excitation wavelengths of the chromophores. Bell-shaped lines display the approximate position of the emission spectra of the different fluorophores and shaded areas indicate the emission filters used to collect photons at different wavelengths. Ratiometric imaging of total GSH concentration using Calnexin-TRaQ-G-mGold (b) and of redox potential (c) using roGFP-iE-ER. The image in (c) is not calibrated and only the ratio of emission at 475 nm upon excitation at either 405 or 445 nm is displayed. d) Comparison of GSH concentrations and 405/445 ratios of roGFP-iE-ER in cells either untreated (ctrl) or after incubation with 10 mM EtGSH for 3 h. *N* = 119, 115, 86, 103 (from left to right) from 3 biological replicates. e) Qualitative monitoring of decrease in GSH concentration using the NIR sensor H2B-TRaQ-G-emiRFP703 (uncalibrated) in cells treated with 1 mM H_2_O_2_ during a period of 20 min. N ≥ 67 cells/timepoint from three biological replicates. Statistical significance was evaluated by unpaired ANOVA except for H_2_O_2_ where paired data were collected. Mean and standard deviation are displayed in panel (d) and individual cell values are depicted in panel (e). ns = P>0.05, * = P≤0.05, ** = P≤0.01, *** = P≤0.001, **** = P≤0.0001.

## 3 Conclusion

We designed, synthesized, and validated a new ratiometric GSH sensor, TRaQ-G, which only becomes reactive in the organelle of interest. This sensor is based on binding of a spirocyclic SiR ligand to HT, which induces isomerization to the GSH-sensitive and fluorescent zwitterionic form. We also created a GSH-insensitive control probe for validation studies. We conducted X-ray diffraction experiments and MD simulations for structural elucidation of the different TRaQ-G adducts to gain mechanistic insight into the varying GSH sensitivities of the molecules. These data could also be used in future studies to improve the kinetics of our sensor by engineering the HT binding pocket, and more generally to design fluorogenic dyes or other robust, targeted sensors. TRaQ-G-mGold was calibrated to measure the absolute GSH concentration of specific subcellular compartments in living cells. When the cellular GSH levels were artificially lowered or raised, TRaQ-G probes reliably detected changes in GSH concentration. We monitored the nuclear and overall GSH levels during cell division for over 24 h. The data suggest an independently regulated nuclear GSH pool and confirm that fluctuations of nuclear GSH are dependent on cell proliferation. TRaQ-G is compatible with roGFP imaging for simultaneous imaging of absolute GSH concentration and organellar redox potential. Our sensor is spectrally flexible, and the fluorescent protein (FP) can be exchanged to any color complementary to the SiR-based ligand. In theory, it would be even possible to exchange the FP for another self-labeling tag to adapt the color for a given experiment. In summary, we presented a robust and flexible tool for live-cell GSH imaging that we used to observe compartmentalization and fluctuations of cellular GSH. We believe that this concept could be extended to develop improved fluorescent sensors for other analytes.

## 4 Methods

### Optical spectroscopic methods

Stock solutions were prepared in DMSO (spectrophotometric grade >99.9%) at concentrations of 50 mM and stored at –20 °C. UV-Vis spectra were acquired using a Multiskan SkyHigh Microplate Spectrophotometer (ThermoFisher Scientific) and 96-well plates (Corning). Fluorescence spectra were acquired using an FS5 Spectrofluorometer (Edinburgh Instruments) equipped with an SC-40 plate reader and 96-well plates (Corning). All measurements were carried out as three technical replicates. The obtained spectra were background corrected. Measurements were carried out at 20 °C unless stated otherwise.

### Cloning

Primers for Gibson assembly were designed using SnapGene. Respective precursor plasmids and primers are reported in Table S1 and Table S2. Reagents and general procedures from the Gibson Assembly Cloning Kit from New England Biolabs (NEB) were used. Vector and insert fragments were linearized by polymerase chain reaction (PCR) and template DNA was digested with DpnI. PCR fragments were purified with the QIAquick PCR Purification Kit from Qiagen. Backbone and insert fragment were ligated and transformed by heat shock into NEB 5α competent cells following the provided standard procedure of the vendor. Plasmids were amplified by incubation of lysogeny broth (LB) cultures containing the appropriate antibiotics overnight at 37 °C. DNA was isolated using the Qiagen Plasmid Mini Kit or the Qiagen Plasmid Plus Midi Kit. The correct sequence of the gene of interest (GOI) was confirmed by the Sanger sequencing service of Microsynth. New constructs reported in this paper are available from Addgene: H2B-HT-emiRFP703 (183985), HT-mGold (183986), Calnexin-HT-mGold (183987), H2B-HT-mGold (183988), and Calnexin-HT8-mGold (183989).

### Site-directed mutagenesis

Primers for site-directed mutagenesis were designed with NEBaseChanger. Respective precursor plasmids and primers are reported in Table S1 and Table S2. The standard procedure and reagents from the Q5 Site-Directed Mutagenesis Kit (NEB) were used to insert mutations, ligate plasmids and transform them into NEB 5α competent cells by heat shock. The correct sequence of the GOI was confirmed by the Sanger sequencing service of Microsynth.

### Protein expression

BL21(DE3) competent cells from NEB were transformed with the respective plasmid by heat shock following the provided standard procedure of the vendor. LB medium was prepared with the appropriate antibiotic. A single colony was inoculated into 100 mL LB medium containing the appropriate antibiotics and incubated at 37 °C overnight. 20 mL of the starter culture were inoculated into 1 L medium and incubated at 37 °C until OD=0.4–0.8 was reached. Isopropyl β-D-1-thiogalactopyranoside (IPTG) was added to a final concentration of 1 mM and the culture was further incubated at 18 °C overnight. *E. coli* were harvested by centrifugation and resuspended in HEPES buffer (20 mM HEPES, 300 mM NaCl, pH=7.4). Glycerol was added to a final concentration of 10% as well as Turbonuclease (5 µL) and a protease inhibitor cocktail tablet (Roche). The cells were lysed by sonication (70% amplitude,10 s pulse/10 s pulse off for 2.5 min). The lysate was cleared by centrifugation and the protein was purified by Ni-His-affinity chromatography in batch-mode. Fractions were analyzed by gel electrophoresis and pure fractions were pooled and dialyzed against PBS.

### Protein crystallization and structural analysis

HT protein (∼1 mg mL^−1^) was reacted with the respective ligand (1.5x excess) in PBS at 20 °C for 1 h. The adduct was purified by size-exclusion chromatography and concentrated to ∼10 mg mL^−1^. The adducts were screened by the sitting-drop vapor diffusion method using commercially available screens from Molecular Dimensions and Qiagen, dispensed by the Mosquito robot (TTP Labtech). Crystals of HT adducts formed in a couple of days in 25% v/v PEG smear medium and 0.1 M Tris pH 8.5 (Me-TRaQ-G), in 20% v/v PEG 6000, 10% ethylene glycol, 0.1 M magnesium chloride hexahydrate and 0.1 M MES pH 6 (TRaQ-G) and in 20% v/v PEG 6000, 0.1 M lithium chloride and 0.1 M sodium citrate pH 4 (TRaQ-G-ctrl). The crystals were cryoprotected with 25% glycerol. Diffraction data were collected at the Paul Scherrer Institute (Swiss light source, Villigen) at the PXIII beamline. Data were processed with the XDS program package^28^. Structures were solved by molecular replacement using Phaser-MR and chain A of PDB entry 6U32 as the model. Manual model building and structure refinement were carried out with PHENIX (version 1.20.1-4487)^29^, Coot (version 0.9.6)^30^ and phenix-refine, respectively. After validation, the structures of the HaloTag adducts were deposited in the PDB database under PDB codes 7ZAA (Me-TRaQ-G), 7ZAB (TRaQ-G), and 7ZAD (TRaQ-G-ctrl). Data collection and refinement statistics are summarized in Table S3. Depictions of adduct structures were generated using PyMOL^31^.

### Molecular dynamics simulations

Atomistic MD simulations were performed starting from the crystal structures of HT with the closed and open form of TRaQ-G probe, as well as with Me-TRaQ-G and TRaQ-G-ctrl probe. All MD simulations were performed with the program package GROMACS (version 2020.4)^32^ using the GROMOS 54a7 force field^33^. The ligand parameters were obtained from the Automated Topology Builder^34,35^.

The ligands were bound to D106 according to the crystal structures using the [intermolecular_interactions] block of GROMACS with a bond length of 0.185 nm, a force constant of 10^5^ kJ mol^−1^ nm^−2^ and bond type 6. The proteins were neutralized and solvated in 0.15 M NaCl solution using the SPC water model and a box size of 7.0 × 7.0 × 7.0 nm^3^. After a steepest decent minimization (5,000 steps), the systems were equilibrated in several steps (1. 500 ps NVT simulation with time step of Δ*t* = 0.5 fs time step; 2. 1 ns NPT simulation with Δ*t* = 1 fs; 3. 1 ns NPT simulation with Δ*t* = 1 fs; 4. 1 ns NPT simulation Δ*t* = 2 fs). During the first two equilibration steps, position restraints were applied to the protein. The production simulations were performed for 500 ns using a time step of Δ*t* = 2 fs. The temperature was kept at 300 K in all simulations using a velocity rescaling thermostat^36^ and the pressure was kept at 1 bar (using a Berendsen barostat^37^ for the equilibration steps and a Parrinello-Rahman barostat^38^ for the production). All bond lengths were constraint using the LINCS algorithm^39^. Van der Waals interactions were treated with a cutoff scheme; Coulomb interactions were calculated using PME. The analyses were performed using GROMACS tools.

In addition, simulations of the GSH adducts of TRaQ-G and Me-TRaQ-G were performed following the protocol described above. The starting conformations were obtained by manually binding GS(H) to a snapshot of the MD simulation of TRaQ-G and the crystal structure of Me-TRaQ-G, respectively.

### *In vitro* GSH sensitivity and selectivity experiments

The HT adducts were prepared by combining the HT protein (∼50 µM in PBS) with the respective TRaQ-G ligand (1.5–2x excess, stock solution 50 mM in DMSO) and rotating the resulting solution for 1 h at 20 °C. The adduct was further concentrated and excess TRaQ-G ligand was removed by desalting with Zeba spin desalting columns (7K MWCO, 0.5 mL). The product was diluted with 0.5 M sodium phosphate buffer to a final concentration of ∼5–15 µM. In a 96-well plate, the adduct solution was treated with the appropriate amount of an aqueous stock solution of GSH, GSSG, cysteine, H_2_S, or taurine. Absorbance at 600 nm and/or fluorescence at 615 nm was measured in triplicates and including separate blanks for every concentration of the reagents.

### Kinetics measurements

HT adducts were prepared as described before. A final concentration of ∼15 µM adduct was used for the measurements. In a 96-well plate, GSH was added (final concentration 5 mM) to the adduct solution and the measurement loop was started. Absorbance was measured in triplicates including a blank measurement without adduct present. After equilibration was reached, *N*-ethylmaleimide (NEM, final concentration 40 mM) or iodoacetamide (final concentration 50 mM) were added and a second measurement loop was started. For Me-TRaQ-G the measurement was carried out at 20 °C, for TRaQ-G the measurement was carried out at 37 °C.

### Calibration curve

The TRaQ-G sensor was assembled *in vitro* as described before using the purified HT-mGold fusion protein. The adduct was used in a final concentration of 10 µM. In a 96-well plate, the adduct was treated with the appropriate amount of GSH. The mGold/SiR ratio was measured by fluorescence microscopy. The ratiometric measurement was carried out at the same instrument with the same settings as for all other measurements. The measurement was performed in three technical replicates and the blank averaged over all concentrations. Three fields of view (FOVs) were measured per well. Image analysis was performed with Fiji (ImageJ). The background was determined from the blank measurements calculating the mean of all the pixels in the FOV per channel excluding the edges. The background was subtracted for the sample measurements and each channel. The mGold channel was divided by the SiR channel in a pixel-by-pixel manner and the mean of all pixels in the FOV excluding the edges was calculated. Every FOV gave one data point.

### Cell culture

HeLa cells were grown in Dulbecco’s Modified Eagle Medium (DMEM) supplemented with fetal bovine serum (FBS, 10%) and penicillin (100 U mL^−1^)/streptomycin (100 µg mL^−1^)/fungizone (0.25 µg mL^−1^) at 37 ºC in 5% CO_2_ environment. For imaging, 15’000–20’000 cells were seeded per well of an 8-well Ibidi chambered cover glass 2–3 days prior to imaging. If required, cells were transfected with plasmid DNA using jetPRIME according to the recommended protocol of the supplier 1–2 days prior to imaging. The cells were incubated with the respective probes in growth medium for the indicated time. Before imaging, the growth medium was removed, the cells were washed with PBS (2x) and imaged in FluoroBrite DMEM.

### Confocal microscopy

Confocal imaging was performed with a dual-camera Nikon W1 spinning disc microscope equipped with an sCMOS camera (Photometrix). Brightfield imaging was performed with a white LED. Laser lines and filters were set up for the appropriate channels as described in (Table S4). Images were collected using a CFI Plan Apochromat Lambda D oil immersion objective (60x, NA = 1.4) and for roGFP imaging a CFI Plan Apo VC water immersion objective (60x, NA = 1.2). Channels were imaged sequentially. The microscope was operated using the NIS elements software. Imaging was performed at 37 ºC in a 5% CO_2_ environment.

### Image analysis

The background was determined by manually picking regions without cells or with untransfected cells. The mean background value for each channel was further used in automated image analysis. Ratiometric image analysis was conducted using a python script (available under https://gitlab.uzh.ch/locbp/public/ratiometric-image-analysis). The background was subtracted for each channel and the regions of interest (= transfected cells) were determined by thresholding or with the cellpose^1^ algorithm in the channel of the FP. To obtain ratiometric images the masks were applied to the background-subtracted images and the FP channel was divided by the SiR channel in a pixel-by-pixel manner within the region of interests (ROIs). Single cells were either identified by selecting coherent regions in a certain range of size or by using the ROIs earlier defined by cellpose. The mean of all ratiometric pixels per cell was calculated and represents one data point. Cells from all replicates were combined for further analyses. The statistical analyses were performed using Prism9 GraphPad and generally included a ROUT outlier analysis. Calculated values are always given as mean ± standard deviation. Our calibration curve was then used to interpolate means as well as upper and lower bounds expressed in GSH concentration. Additionally, our linear calibration equation was applied to the ratiometric images and saved for each FOV. For display only, images were despeckled with Fiji (ImageJ).

### GSH measurement in living HeLa cells with external manipulation of the GSH concentration

HeLa cells were plated and transfected with the respective HT-mGold plasmid. After 24–48 h the cells were incubated with 100 nM TRaQ-G ligand for 1 h. The cells were treated with 10 mM EtGSH in FluoroBrite DMEM for 3–4 h, 1 mM BSO in FluoroBrite DMEM for 3–4 h, 1 mM H_2_O_2_ for 20 min in FluoroBrite or vehicle (FluoroBrite DMEM). GSH concentration was measured by fluorescence microscopy using the calibration curve. The experiment was carried out in three biological replicates on different days with cells from different passages. The data sets were combined for analysis. In total, 90–176 cells were analyzed per condition.

### GSH measurement during cell proliferation

HeLa cells were plated and transfected with the HT-mGold or H2B-HT-mGold plasmid. About 8 h after transfection, thymidine (100 mM aqueous stock solution, final concentration 2 mM) was added to the cells and incubated for 16 h. The cells were washed with PBS and incubated in fresh medium for 9 h. Thymidine was added again (final concentration 2 mM) and incubated for 18 h. Before imaging, the cells were incubated with 100 nM TRaQ-G or TRaQ-G-ctrl ligand for 1 h in culture medium still containing thymidine. Cells were imaged in fresh FluoroBrite DMEM supplemented with 4 mM glutamine, 1 mM sodium pyruvate, 10% FBS and penicillin (100 U mL^−1^)/streptomycin (100 µg mL^−1^)/fungizone (0.25 µg mL^−1^). Frames were taken every 30 min over 24 h. For analysis, ratiometric images of the whole FOV were generated with the method described before. From each FOV the dividing cells were segmented and followed over time by hand. The mean of all pixels within one cell represents one data point. The experiment was carried out in three biological replicates on different days with cells from different passages. The data sets were combined for analysis. In total, a minimum of 19 cells were analyzed per condition and frame.

### Color imaging with TRaQ-G and roGFP

HeLa cells were plated and co-transfected with ER-Halo-mGold and roGFP-iE-ER (3:1). After 24–48 h, cells were incubated with the TRaQ-G ligand for 1 h. Cells were treated with 10 mM EtGSH in FluoroBrite DMEM for 3–4 h or vehicle (FluoroBrite DMEM). The same cells were imaged with the 60x oil immersion objective for the GSH concentration and the 60x water immersion objective for the redox potential. The experiment was carried out in three biological replicates on different days with cells from different passages. The data sets were combined for analysis. In total, 86–119 cells were imaged per condition.

## 5 Extended Data

**Extended Data Figure 1.**
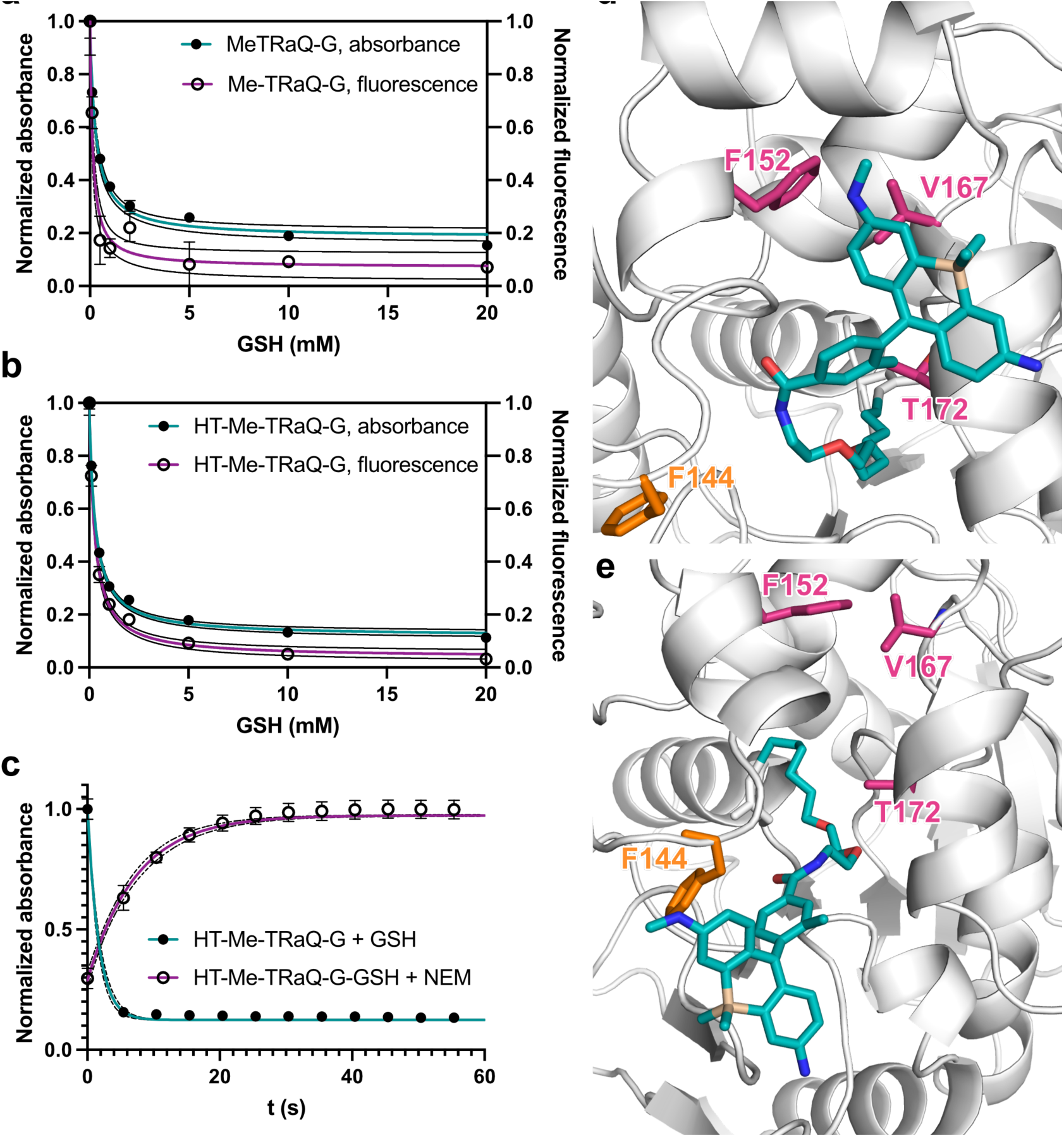
a) Me-TRaQ-G shows good GSH-sensitivity in the range of 0–5 mM by itself and b) as HT adduct. c) The conjugate with GSH forms rapidly in vitro and is reversible by adding the GSH scavenger NEM. Data were fitted by one-phase decay, dotted lines represent 95% confidence interval (CI). d) Preferred conformation of Me-TRaQ-G during MD simulation (322 ns) with methyl group pointing into hydrophobic pocket. e) Short-lived conformation of Me-TRaQ-G during MD simulation (457 ns) with methyl group exposed to the solvent. Residues forming the hydrophobic pocket are displayed in pink, the Phe residue that moves between the open and closed conformation is displayed in orange.

**Extended Data Figure 2.**
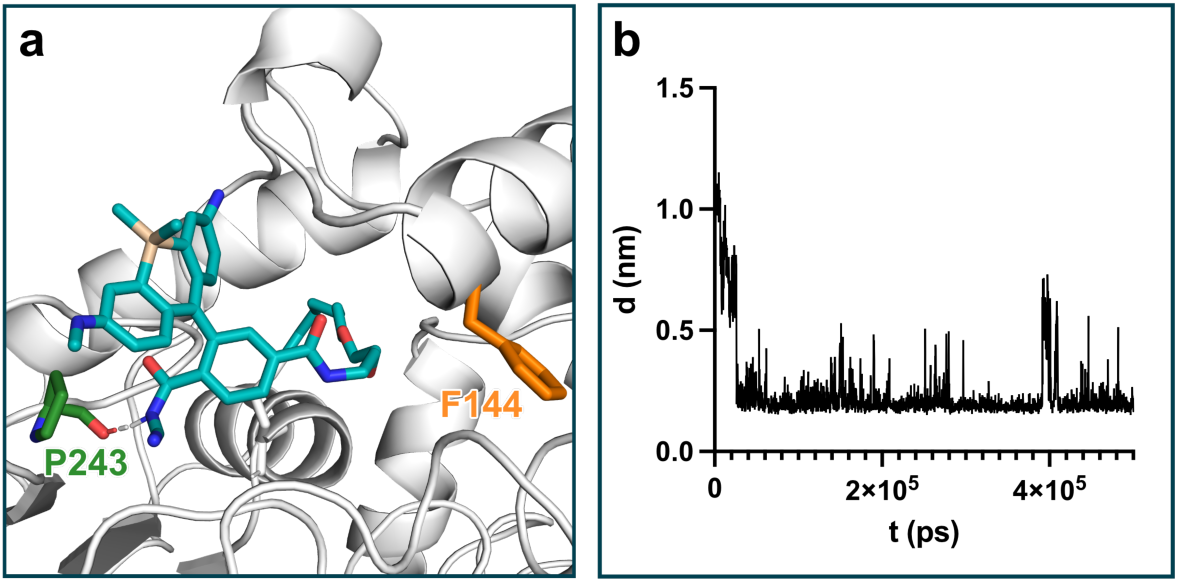
a) Snapshot during MD simulation of open TRaQ-G at 102 ns. b) Length of the hydrogen bond between the cyanamide proton and the backbone carbonyl of P243 during the simulation. The Phe residue that moves between the open and closed conformation is displayed in orange and the residue forming a hydrogen bond with the ligand is displayed in green.

**Extended Data Figure 3.**
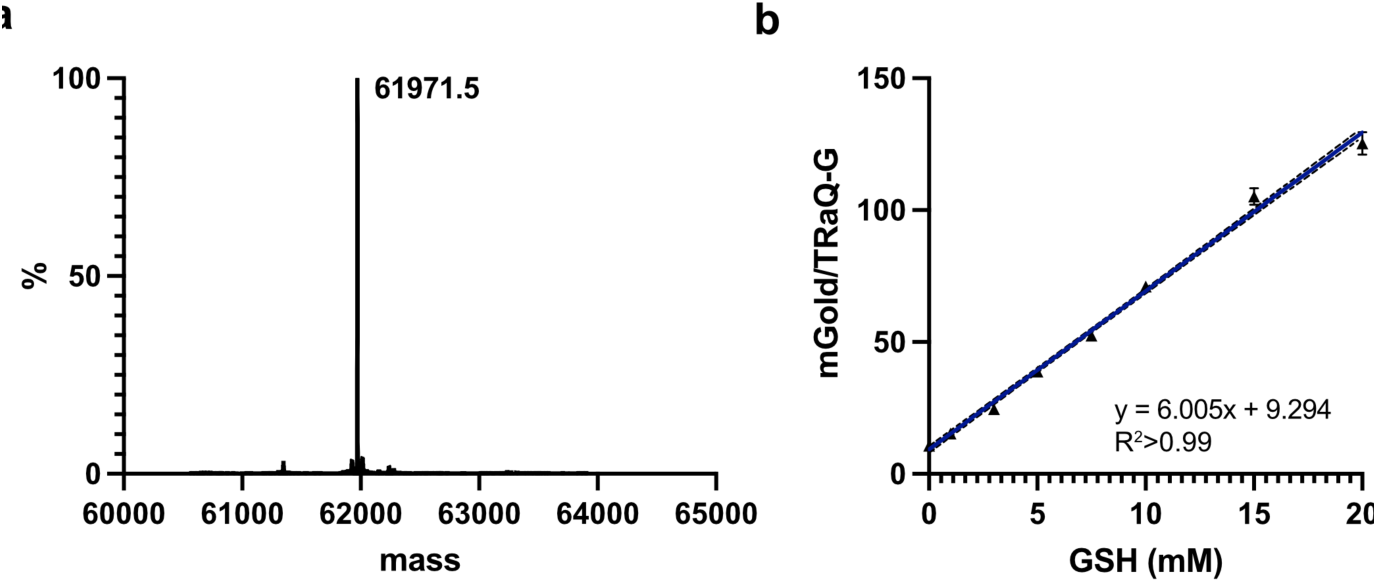
a) Deconvoluted mass spectrum of TRaQ-G reacted with HT *in vitro*. b) Calibration curve of TRaQ-G with adduct formed by *in vitro* reaction of purified fusion protein with ligand and measured by fluorescence microscopy. Data were fitted by linear regression. Dashed line represents 95% CI.

## Acknowledgements

This work was funded by the Swiss National Science Foundation (grant PCEGP2_186862). We thank the whole PTPSP team at EPFL, especially Florence Pojer, Kelvin Lau and Amédé Larabi for their help with protein expression, purification, crystallization, and X-ray diffraction analyses. X-ray diffraction data were collected at the PXIII beamline of the Swiss Light Source, Paul Scherrer Institute (Villigen, Switzerland). S. T. thanks the Alfons und Gertrud Kassel foundation, the Dr. Rolf M. Schwiete foundation and the Center for Multiscale Modelling in Life Sciences (CMMS) sponsored by the Hessian Ministry of Science and Art for funding and the Center for Scientific Computing at the Goethe University Frankfurt for access to Goethe-HLR.

## Author contributions

S. H. and P. R.-F. conceived the project. S. H. performed synthesis of small molecules, cloning of plasmids, protein expression and purification, titrations, kinetics experiments, X-ray diffraction analyses, cell culture, imaging, data analysis, and manuscript preparation. G. Q. contributed to the synthesis of TRaQ-G-ctrl, to cloning of some plasmids, and to image acquisition and analysis. S. T. performed and analyzed MD simulations. P. R.-F. supervised the project and contributed to data analysis and manuscript preparation.

## Competing interests

The authors declare no competing interests.

## Notes

### Competing Interest Statement

The authors have declared no competing interest.

