## Supporting Information for "A Locally Activatable Sensor for Robust Quantification of Organellar Glutathione"

### 1 General remarks

Unless stated otherwise, all reagents and solvents were purchased from commercial sources and used as received. NMR spectra were acquired on Bruker AVANCE NEO-400, Bruker AVANCE III-400, Bruker AVANCE III HD-600, and Bruker AVANCE II-800 instruments. $^{1}$H NMR chemical shifts are reported in ppm relative to $SiMe_4$ ($\delta$ = 0) and were referenced internally with respect to residual protons in the solvent ($\delta$ = 1.94 for acetonitrile and $\delta$ = 3.31 for methanol)[1]. Coupling constants are reported in Hz. $^{13}$C NMR chemical shifts are reported in ppm relative to $SiMe_4$ ($\delta$ = 0) and were referenced internally with respect to solvent signal ($\delta$ = 1.32 for acetonitrile and $\delta$ = 49.00 for methanol)[1]. High-resolution mass spectrometry (HRMS) was performed by the MS facility of EPFL. Ultra-high performance liquid chromatography – mass spectrometry (UHPLC-MS) analysis of the ligands in the presence of GSH was performed on a Waters system with an ACQUITY UHPLC and a SYNAPT-G2 mass spectrometer using electrospray ionization (ESI). Reaction progress was followed by thin-layer chromatography (TLC) and UHPLC-MS on a Shimadzu LC-MS 2020 system using ESI. Purification by flash column chromatography and prep-HPLC was performed using a Büchi Pure-Chromatography-System and Büchi FlashPure columns. IUPAC names of all compounds are provided and were determined using CS ChemDraw 19.1.

#### 2 Supplementary Figures

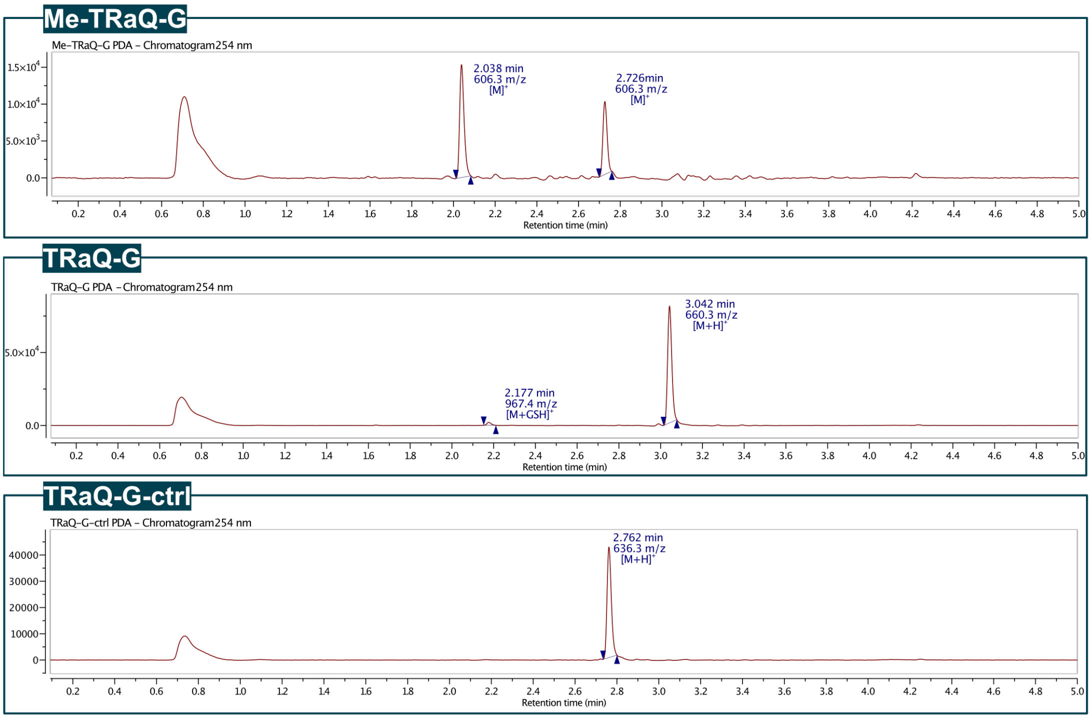

**Figure S1**. LC-MS analysis of Me-TRaQ-G, TRaQ-G and TRaQ-G-ctrl at 20 µM with 10 mM GSH added. Gradient 10 → 95% MeCN in dd$H_2O$ + 0.02% trifluoro-acetic acid (TFA) and 0.04% formic acid. Me-TRaQ-G adduct with GSH (2.038 min) is not stable during ionization and is therefore detected as fragment with loss of GSH. TRaQ-G shows traces of GSH adduct after 10 min. The amount of formed adduct increases slowly with incubation time. The impact on live cell experiments should be minimal as incubation times can be kept short (≤1 h) and diffusion of the GSH adduct is expected to be much slower than for the ligand itself. TRaQ-G-ctrl is not glutathionylated even after ~2 h incubation.

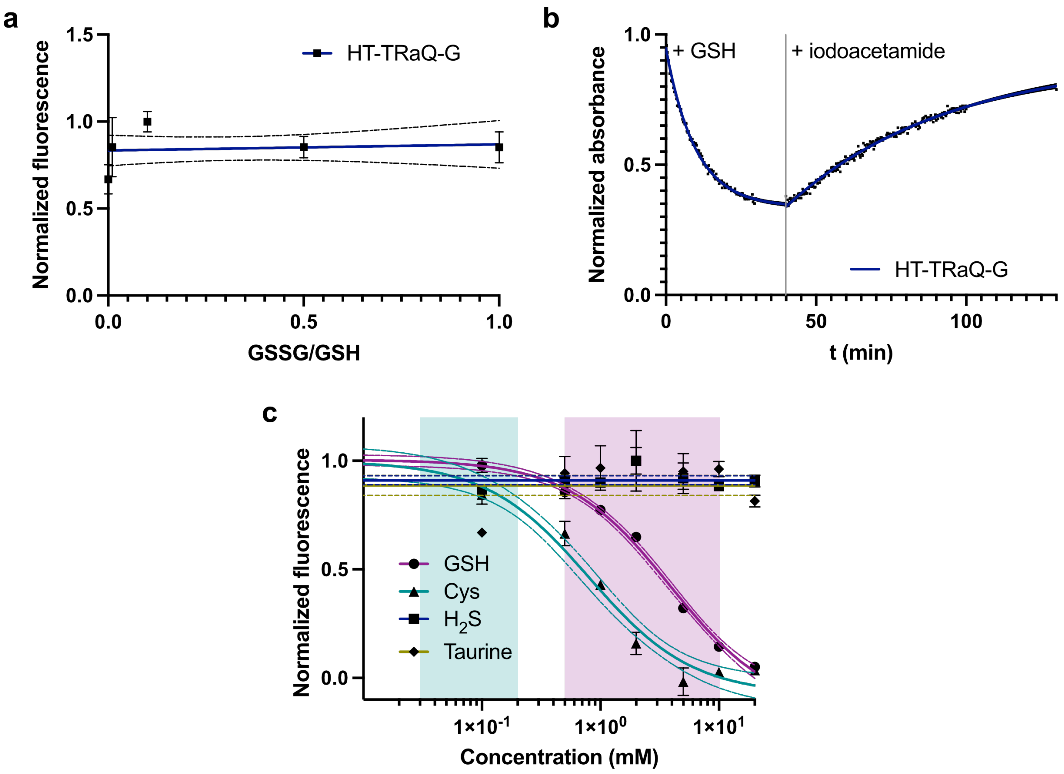

**Figure S2**. a) Sensitivity of HT-TRaQ-G to the ratio between oxidized and reduced glutathione. Data were fitted by linear regression. Dotted lines represent the 95% CI. b) Time-resolved response of HT-TRaQ-G to GSH and reversibility after the addition of the thiol-scavenger iodoacetamide. c) GSH-sensitivity of HT-TRaQ-G compared to its sensitivity towards cysteine, $H_2S$ and taurine. Range of intracellular concentration is depicted in magenta for GSH[2] and cyan for cysteine[3]. Data were fitted by dose-response model (GSH, cysteine) or by linear regression ($H_2S$, taurine). Dotted lines represent the 95% CI.

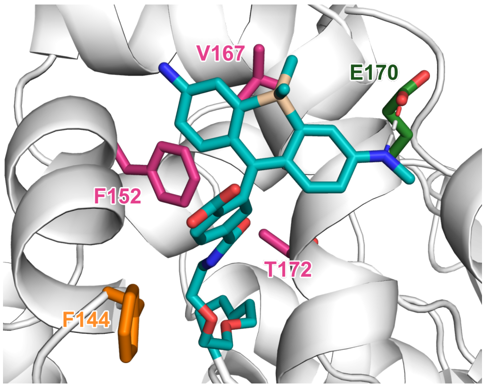

**Figure S3**. Snapshot of HT-TRaQ-G-ctrl during MD simulation (444 ns) forming a hydrogen bond to Glu170 and exposing the carboxylate group to the solvent. Residues forming the hydrophobic pocket are displayed in pink, the Phe residue that moves between the open and closed conformation is displayed in orange and the residue forming a hydrogen bond with the ligand is displayed in green.

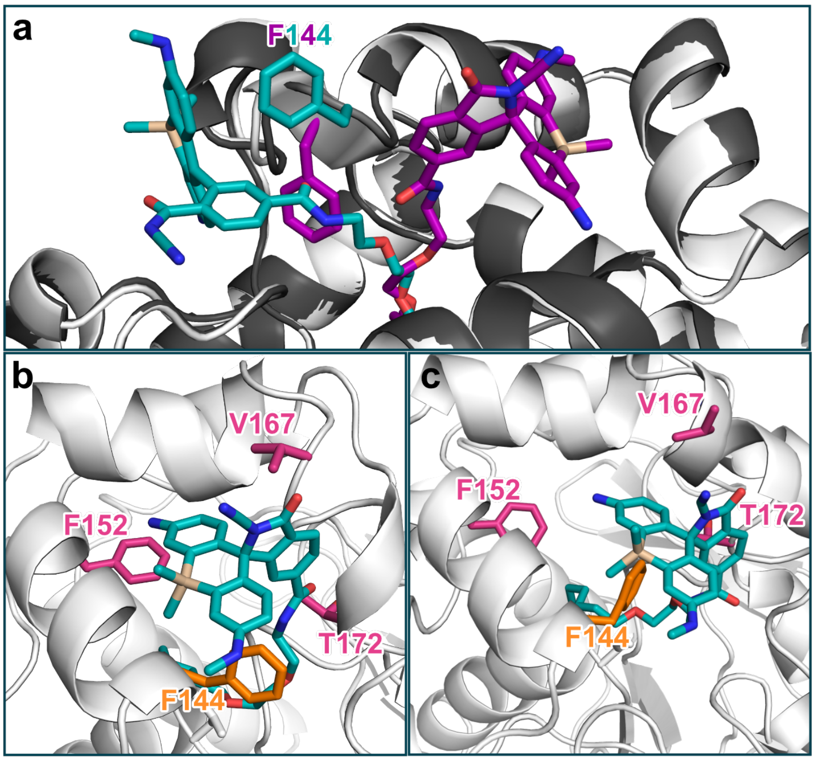

**Figure S4**. a) Overlaid X-ray diffraction structures of the two chains in the TRaQ-G crystal with the closed form in magenta/white and the open form in cyan/gray. The F144 residue is displayed in its respective conformation with color corresponding to the TRaQ-G ligand of the same chain. b) Snapshot of MD simulation of closed TRaQ-G at 167 ns in its original position and at 216 ns when crossed F144. Residues forming the hydrophobic pocket are displayed in pink, the Phe residue that moves between the open and closed conformation is displayed in orange or in the color of the respective conformation in panel (a).

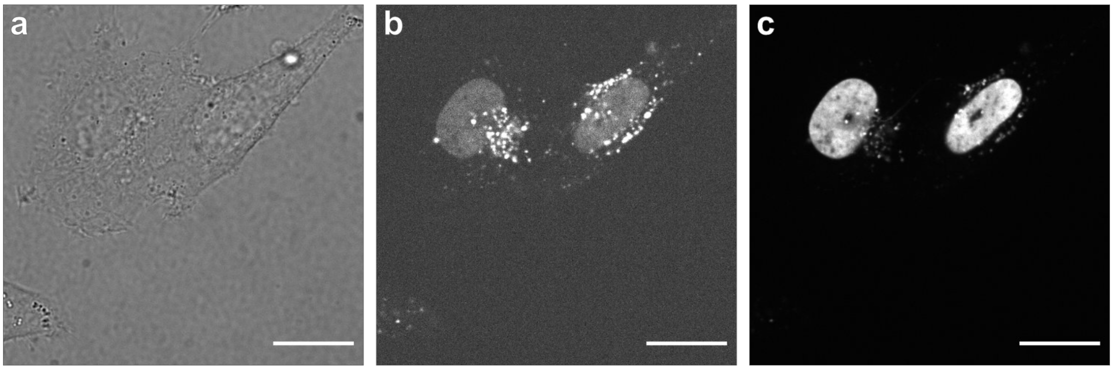

**Figure S5**. Imaging of Me-TRaQ-G in HeLa cells expressing H2B-HT-emiRFP703 in the a) brightfield, b) Me-TRaQ-G and c) emiRFP703 channel. Scale bars = 20 µm.

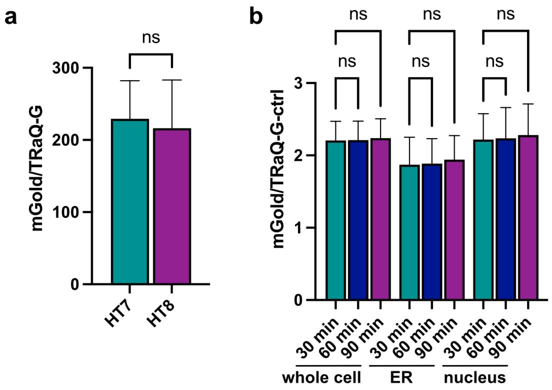

**Figure S6**. a) Comparison of HT7 to redox-insensitive HT8 displaying similar labeling efficiency with TRaQ-G. *N* = 37 (HT7) and 30 (HT8) from one biological replicate. b) Saturation of the HT-mGold fusion protein with TRaQ-G-ctrl in living HeLa cells. *N* = 90, 91, 91, 142, 131, 101, 116, 115, 111 (from left to right) from 3 biological replicates. ns = P>0.05, * = P≤0.05, ** = P≤0.01, *** = P≤0.001, **** = P≤0.0001.

### 3 Supplementary Tables

**Table S1.** Plasmid sources.

| Plasmid | Source | Vector precursor | Insert precursor |
|---|---|---|---|
| Calnexin-mGold | Addgene 158004[4] | | |
| TUBB5-Halo | Addgene 64691[5] | | |
| H2B-emiRFP703 | Addgene 136567[6] | H2B-emiRFP703 | TUBB5-Halo |
| H2B-HT-emiRFP703 | Gibson assembly | | |
| HT-mGold | Gibson assembly | Calnexin-mGold | TUBB5-Halo |
| ER-HT-mGold | Gibson assembly | Calnexin-mGold | TUBB5-Halo |
| H2B-HT-mGold | Gibson assembly | H2B-HT-emiRFP703 | ER-HT-mGold |
| ER-HT8-mGold | Site-directed mutagenesis | ER-HT-mGold | |
| HaloTag7-His6 | Gift from Thomas Ward (University of Basel) | | |
| HT-mGold-His6 | Twist Bioscience | | |
| roGFP-iE-ER | Gift from Christian Appenzeller-Herzog (University of Basel). | | |

**Table S2.** Primers for plasmid generation.

| Plasmid | Vector forward | Vector reverse | Insert forward | Insert reverse |
|---|---|---|---|---|
| H2B-HT-emiRFP703 | atttccggcgatccaccggt | gtcgactgcttagcgctggt | agcgctaagcagtcgaccg | ggtggatcgccggaaatctcg |
| HT-mGold | atttccggcgtgagcaagggc | ccgatttccatggcggtctc | cgccatggaaatcggtactgg | tgctcacgccggaaatctc |
| ER-HT-mGold | atttccggcgtgagcaaggg | tcgactgcatggtggcga | accatgcagtcgaccggca | ttgctcacgccggaaatctcg |
| H2B-HT-mGold | gtacaagtaagagagctaagcggc | tggccattcgtcccaggt | gacctgggacgaatggcca | ccgcttagctctcttacttgtacagctc |
| ER-HT8-mGold C61S | gacccatcgcagcattgctcc | ggtgcaacatgcgggatg | | |
| ER-HT8-mGold C262S | cctgcctaacagcaaggctgtg | cttttggccaggcgagcg | | |

**Table S3.** Data collection and refinement statistics.

| | Me-TRaQ-G | TRaQ-G | TRaQ-G-ctrl |
|---|---|---|---|
| **Data collection** | | | |
| Space group | P 21 21 21 | C 1 2 1 | P 61 |
| Cell dimensions | | | |
| *a*, *b*, *c* (Å) | 44.36, 81.50, 159.65 | 166.91, 50.46, 79.48 | 94.01, 94,01, 132.56 |
| α, β, γ (°) | 90, 90, 90 | 90, 117.737, 90 | 90, 90, 120 |
| Resolution (Å) | 1.23–50 (1.23–1.3) | 1.68–50 (1.68–1.79) | 1.95–44.3 (1.95–2.06) |
| $R_{meas}$ (%) | 8.8 (153.6) | 11.1 (109.2) | 20.7 (114.6) |
| $I$ / σ$I$ | 14.16 (1.15) | 7.24 (0.98) | 6.42 (1.55) |
| Completeness (%) | 99.8 (98.7) | 98.4 (96.2) | 99.8 (99.2) |
| Redundancy | 10.5 (8.2) | 3.0 (2.6) | 4.6 (4.4) |
| **Refinement** | | | |
| Resolution (Å) | 1.23 | 1.68 | 1.95 |
| No. reflections | 168606 | 65995 | 94488 |
| $R_{\text{work}}$ / $R_{\text{free}}$ | 0.23 / 0.25 | 0.19 / 0.22 | 0.18 / 0.22 |
| No. atoms | | | |
| Protein | 4699 | 4694 | 4730 |
| Ligand/ion | 294 | 205 | 216 |
| Water | 478 | 434 | 436 |
| *B*-factors | | | |
| Protein | 16.94 | 26.40 | 22.61 |
| Ligand/ion | 33.29 | 29.84 | 30.34 |
| Water | 26.60 | 33.94 | 29.81 |
| R.m.s. deviations | | | |
| Bond lengths (Å) | 0.005 | 0.012 | 0.008 |
| Bond angles (°) | 0.865 | 1.173 | 1.004 |

**Table S4.** Microscope settings for imaging channels.

| Channel | λ excitation | Emission filter |
|---|---|---|
| roGFP blue | 405 nm | 472/30 |
| roGFP green | 445 nm | 472/30 |
| mGold | 515 nm | 542/27 |
| SiR with mGold | 561 nm | 642 LP |
| SiR with emiRFP703 | 561 nm | 600/52 |
| emiRFP703 | 638 nm | 708/75 |

### 4 Synthetic procedures

#### 4.1 Synthesis of the Me-TRaQ-G ligand

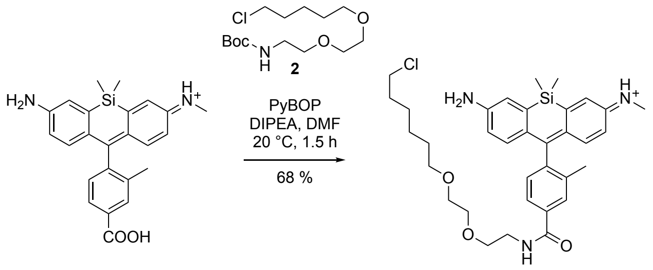

1    Me-TRaQ-G

**Scheme S1**. Synthesis of the **Me-TRaQ-G** ligand.

Intermediate **1** was prepared according to previously reported procedures[3].

(*E*)-*N*-(7-Amino-10-(4-((2-(2-((6-chlorohexyl)oxy)ethoxy)ethyl)carbamoyl)-2-methylphenyl)-5,5-dimethyldibenzo[*b*,*e*]silin-3(5*H*)-ylidene)methanaminium (**Me-TRaQ-G**)

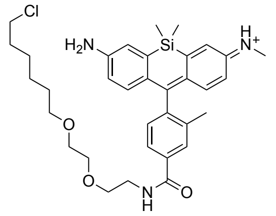

Intermediate **1** (15.0 mg, 29.2 µmol, 1 equiv.), chloroalkane **2** (32.6 mg, 146 µmol, 5 equiv.), and PyBOP (30.3 mg, 58.3 µmol, 2 equiv.) were added to a pre-dried flask under an $N_2$ atmosphere. The solids were dissolved in *N*,*N*-dimethylformamide (1.5 mL) and treated with diisopropylethylamine (DIPEA, 72 µL, 0.44 µmol, 15 equiv.). The resulting solution was stirred at 20 °C for 1.5 h. All volatile material was evaporated under reduced pressure, and the residue was purified by preparative HPLC (10 → 90 % MeCN in dd$H_2O$ + 0.1% TFA over 25 min) to yield a blue solid (13.9 mg, 19.7 µmol, 68%).

$^1$H NMR (400 MHz, MeOD) *δ* 7.54 (d, *J* = 7.7 Hz, 1H), 7.22 – 7.12 (m, 5H), 7.08 (d, *J* = 9.3 Hz, 1H), 6.63 – 6.58 (m, 1H), 6.56 (dd, *J* = 9.3, 2.5 Hz, 1H), 3.73 – 3.66 (m, 4H), 3.63 (m, 4H), 3.50 (m, 4H), 3.07 (s, 3H), 2.49 (s, 3H), 1.72 (dt, *J* = 8.0, 6.5 Hz, 2H), 1.59 (p, *J* = 6.8 Hz, 2H), 1.48 – 1.34 (m, 2H), 0.55 (s, 6H).

$^{13}$C NMR (201 MHz, MeOD) *δ* 172.38, 170.61, 162.48, 158.20, 157.39, 150.21, 144.28, 142.07, 138.37, 137.20, 133.50, 132.39, 130.16, 128.79, 128.75, 128.00, 127.95, 127.77, 124.36, 116.46, 72.25, 71.27, 71.26, 70.45, 45.69, 45.67, 40.80, 33.73, 30.59, 27.78, 26.50, 19.73, -1.42, -1.47.

HRMS (ESI/LTQ-Orbitrap) $[M+H]^+$ calculated for $C_{34}H_{45}ClN_3O_3Si^+$ 606.2913; found 606.2927.

#### 4.2 Synthesis of the TRaQ-G ligand

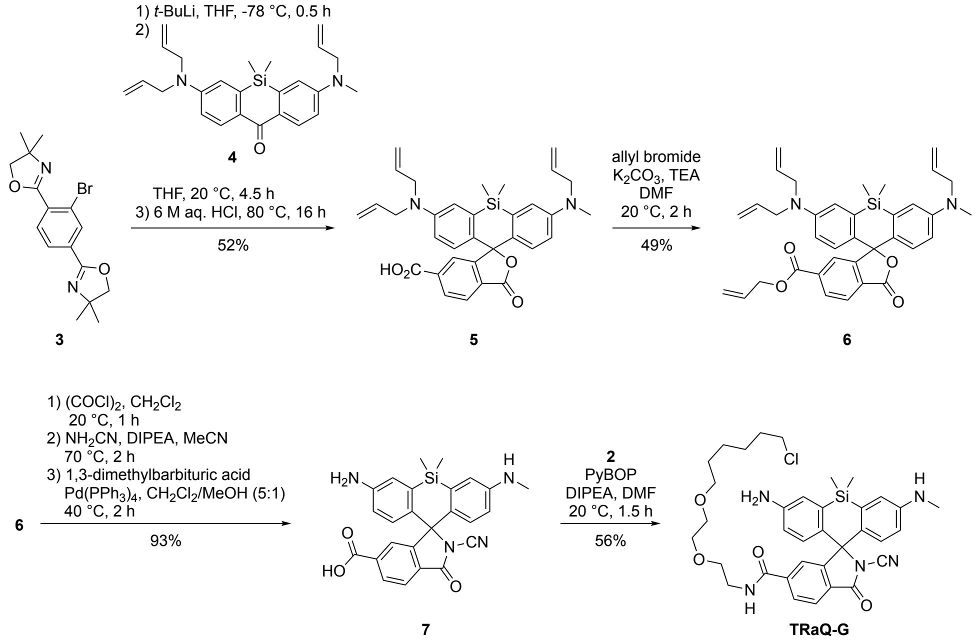

**Scheme S2**. Synthesis of the **TRaQ-G** ligand.

Intermediates **3** and **4** were prepared according to reported procedures[3,7].

3-(Allyl(methyl)amino)-7-(diallylamino)-5,5-dimethyl-3'-oxo-3'*H*,5*H*-spiro[dibenzo[*b,e*]siline-10,1'-isobenzofuran]-6'-carboxylic acid (**5**)

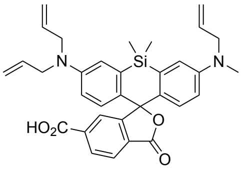

Compound **3** (650 mg, 1.85 mmol, 3 equiv.) was added to a pre-dried flask under an $N_2$ atmosphere, dissolved in dry tetrahydrofuran (31 mL), and cooled to –78 °C. *tert*-Butyllithium (1.7 M, 1.1 mL, 1.9 mmol, 3 equiv.) was slowly added dropwise and the solution was stirred at the same temperature for 30 min. Ketone **4** (249 mg, 617 µmol, 1 equiv.) in dry tetrahydrofuran (15 mL) was added dropwise via a syringe, and the solution was warmed to ambient temperature and stirred for 4.5 h. Acetic acid (3.5 mL, 62 mmol, 100 equiv.) was added to the mixture and the resulting intensely blue solution was evaporated under reduced pressure. The crude product was dissolved in hydrochloric acid (6 M, 51 mL, 500 equiv.) and stirred at 80 °C for 16 h. After cooling to ambient temperature, the solution was added to saturated aqueous $Na_2CO_3$ (50 mL), the pH was adjusted to ~2 and the mixture was extracted with $CH_2Cl_2$ (3x). The combined organic phases were dried over $Na_2SO_4$ and evaporated under reduced pressure. The residue was purified by flash column chromatography ($SiO_2$, 0 → 5% MeOH in $CH_2Cl_2$) to yield a green solid (178 mg, 322 µmol, 52%).

$^1$H NMR (800 MHz, MeOD) δ 8.27 (dd, *J* = 8.1, 1.5 Hz, 1H), 8.18 (d, *J* = 8.1 Hz, 1H), 7.84 (s, 1H), 7.18 (d, *J* = 30.2 Hz, 2H), 6.84 (dd, *J* = 26.2, 9.3 Hz, 2H), 6.70

(dd, *J* = 22.3, 9.1 Hz, 2H), 5.93 – 5.84 (m, 3H), 5.24 – 5.13 (m, 6H), 4.14 (d, *J* = 18.0 Hz, 6H), 3.15 (s, 3H), 0.61 (s, 3H), 0.53 (s, 3H).
$^{13}$C NMR (201 MHz, MeOD) *δ* 172.35, 168.02, 162.65, 150.50, 149.51, 136.53, 136.46, 136.42, 133.87, 133.41, 131.26, 131.21, 130.88, 129.78, 129.58, 126.30, 120.27, 120.16, 117.41, 117.27, 116.48, 115.22, 115.16, 100.02, 56.02, 54.20, 38.99, -0.42, -1.69.
HRMS (ESI/QTOF) $[M+H]^+$ calculated for $[C_{33}H_{35}N_2O_4Si]^+$ 551.2361; found 551.2373.

Allyl 3-(allyl(methyl)amino)-7-(diallylamino)-5,5-dimethyl-3'-oxo-3'*H*,5*H*-spiro-[dibenzo[*b*,*e*]siline-10,1'-isobenzofuran]-6'-carboxylate (**6**)

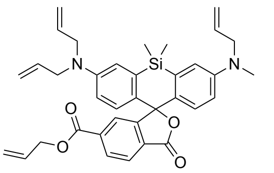

Compound **5** (150 mg, 272 µmol, 1 equiv.) was added to a pre-dried flask under an $N_2$ atmosphere, dissolved in *N*,*N*-dimethylformamide (3.9 mL) and treated with potassium carbonate (75.1 mg, 544 µmol, 2 equiv.) and triethylamine (75.6 µL, 544 µmol, 2 equiv. The mixture was cooled with an ice bath and allyl bromide (35.5 µL, 408 µmol, 1.5 equiv.) was slowly added. The reaction was stirred at 20 °C for 2 h. The resulting solution was diluted with water and extracted with $CH_2Cl_2$ (3x). The combined organic layers were dried over $Na_2SO_4$ and concentrated under reduced pressure. The crude product was purified by flash column chromatography ($SiO_2$, 0 → 25 % EtOAc in hexane) to yield a yellow solid (78 .0 mg, 132 µmol, 49%).
$^1$H NMR (600 MHz, $CD_3CN$) *δ* 8.19 (dd, *J* = 8.1, 1.4 Hz, 1H), 8.01 (d, *J* = 8.0 Hz, 1H), 7.79 – 7.76 (m, 1H), 7.05 (dd, *J* = 15.2, 2.9 Hz, 2H), 6.73 (dd, *J* = 16.4, 9.0 Hz, 2H), 6.59 (ddd, *J* = 19.2, 9.0, 2.9 Hz, 2H), 6.01 (ddt, *J* = 17.2, 10.4, 5.7 Hz, 1H), 5.90 – 5.78 (m, 3H), 5.36 (dq, *J* = 17.3, 1.6 Hz, 1H), 5.25 (dq, *J* = 10.5, 1.4 Hz, 1H), 5.17 – 5.07 (m, 6H), 4.76 (dt, *J* = 5.8, 1.4 Hz, 2H), 4.00 – 3.95 (m, 6H), 2.97 (s, 3H), 0.60 (s, 3H), 0.51 (s, 3H).
$^{13}$C NMR (151 MHz, $CD_3CN$) *δ* 170.10, 165.63, 155.23, 149.67, 148.88, 137.99, 137.84, 136.49, 134.78, 134.30, 133.13, 131.54, 131.36, 130.86, 130.71, 129.79, 129.70, 127.15, 126.02, 119.09, 118.12, 118.05, 116.77, 116.54, 114.78, 114.69, 67.05, 55.33, 53.48, 38.58, -0.05, -1.17. (One carbon is likely hidden under the solvent peak.)
HRMS (ESI/QTOF) $[M+H]^+$ calculated for $C_{36}H_{39}N_2O_4Si^+$ 591.2674; found 591.2687.

3-Amino-2'-cyano-5,5-dimethyl-7-(methylamino)-3'-oxo-5*H*-spiro[dibenzo[*b*,*e*]-siline-10,1'-isoindoline]-6'-carboxylic acid (**7**)

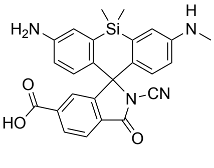

Compound **6** (70.0 mg, 118 µmol, 1 equiv.) was added to a pre-dried flask under an $N_2$ atmosphere, dissolved in dry $CH_2Cl_2$ (20 mM, 5.9 mL) and treated with oxalyl chloride (0.50 M, 0.36 mL, 1.5 equiv.) at 0 °C. The resulting solution was stirred at 20 °C for 2 h. The solvent was evaporated under reduced pressure and the product was used without further purification for the next step.

The crude acyl chloride was dissolved in dry acetonitrile (5.9 mL) under an $N_2$ atmosphere and treated with a solution of cyanamide (49.8 mg, 1.18 mmol, 10 equiv.) and DIPEA (294 µL, 1.78 mmol, 15 equiv.) in dry acetonitrile (3.0 mL). The resulting solution was stirred at 70 °C for 3 h. The solvent was evaporated under reduced pressure and the product was used without further purification in the next step.

The crude cyanamide, 1,3-dimethyl-1,3-diazinane-2,4,6-trione (370 mg, 2.37 mmol, 20 equiv.) and tetrakis(triphenylphosphine)-palladium(0) (68.5 mg, 59.2 µmol, 0.5 equiv.) were dissolved in a degassed mixture (5:1) of $CH_2Cl_2$ (5.9 mL) and methanol (1.2 mL) under an $N_2$ atmosphere. The resulting mixture was stirred at 40 °C for 2 h. The reaction was diluted with $CH_2Cl_2$ and washed with saturated aqueous $Na_2CO_3$ solution. The aqueous phase was re-extracted with $CH_2Cl_2$ (2x). The combined organic phases were dried over $Na_2SO_4$ and the solvent was evaporated under reduced pressure. The crude product was purified by flash column chromatography ($SiO_2$, 0 → 10 % MeOH in $CH_2Cl_2$) to yield a yellowish solid (50.2 mg, 110 µmol, 93%).

$^1$H NMR (600 MHz, MeOD) *δ* 8.18 (dd, *J* = 8.1, 1.3 Hz, 1H), 8.08 (d, *J* = 8.0 Hz, 1H), 7.55 (s, 1H), 7.37 (d, *J* = 2.6 Hz, 1H), 7.00 (dt, *J* = 4.9, 3.0 Hz, 2H), 6.89 (d, *J* = 8.7 Hz, 1H), 6.75 (d, *J* = 8.8 Hz, 1H), 6.70 (dd, *J* = 8.9, 2.6 Hz, 1H), 2.83 (s, 3H), 0.65 (s, 3H), 0.59 (s, 3H).

$^{13}$C NMR (151 MHz, MeOD) *δ* 168.34, 167.51, 162.48, 156.31, 149.82, 141.93, 139.03, 138.32, 136.51, 131.30, 130.23, 130.11, 129.83, 129.25, 126.27, 126.05, 124.25, 122.23, 118.07, 117.26, 107.51, 75.84, 30.37, 0.02, -0.14.

HRMS (ESI/QTOF) $[M+H]^+$ calculated for $C_{25}H_{23}N_4O_3Si^+$ 455.1534; found 455.1542.

3-Amino-*N*-(2-(2-((6-chlorohexyl)oxy)ethoxy)ethyl)-2'-cyano-5,5-dimethyl-7-(methylamino)-3'-oxo-5*H*-spiro[dibenzo[*b*,*e*]siline-10,1'-isoindoline]-6'-carboxamide (**TraQ-G**)

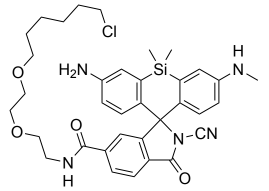

Chloroalkane **2**, compound **7** (20.0 mg, 44.0 µmol, 1 equiv.) and PyBOP (45.8 mg, 88.0 µmol, 2 equiv.) were added to a pre-dried flask under an $N_2$ atmosphere, dissolved in *N*,*N*-dimethylformamide (2.2 mL) and treated with DIPEA (72.7 µL, 440 µmol, 10 equiv.). The resulting solution was stirred at 20 °C for 2.5 h. All volatiles were removed under reduced pressure and the residue was purified by preparative HPLC (10 → 95 % MeCN in dd$H_2O$ + 0.1% TFA over 30 min) to yield a light green solid (16.3 mg, 24.5 mmol, 56%).

$^1$H NMR (600 MHz, $CD_3CN$) *δ* 8.01 (d, *J* = 8.0 Hz, 1H), 7.89 (dd, *J* = 8.0, 1.4 Hz, 1H), 7.29 – 7.27 (m, 1H), 7.19 (s, 1H), 7.06 (t, *J* = 2.0 Hz, 1H), 7.00 (q, *J* = 2.6 Hz, 1H), 6.76 – 6.73 (m, 1H), 6.72 – 6.65 (m, 3H), 3.53 (t, *J* = 6.7 Hz, 2H), 3.51 – 3.48 (m, 4H), 3.45 – 3.37 (m, 4H), 3.30 (t, *J* = 6.5 Hz, 2H), 2.81 (s, 3H),

1.67 (dt, $J$ = 14.7, 6.8 Hz, 2H), 1.43 – 1.37 (m, 2H), 1.36 – 1.30 (m, 2H), 1.26 – 1.19 (m, 2H), 0.58 (s, 3H), 0.52 (s, 3H).
$^{13}$C NMR (151 MHz, $CD_3CN$) $\delta$ 168.04, 166.34, 159.61, 156.21, 148.70, 147.14, 142.85, 137.06, 131.20, 130.81, 130.16, 130.10, 128.72, 127.72, 126.07, 123.58, 120.21, 119.05, 117.87, 117.34, 107.85, 75.49, 71.50, 70.81, 70.68, 69.76, 46.20, 40.50, 33.25, 31.17, 30.16, 27.29, 26.06, -0.04. (Two carbons adjacent to silicon carbons are likely overlapping.)
HRMS (ESI/QTOF) $[M+H]^+$ calculated for $C_{35}H_{43}ClN_5O_4Si^+$ 660.2767; found 660.2787.

#### 4.3 Synthesis of the TRaQ-G-ctrl ligand

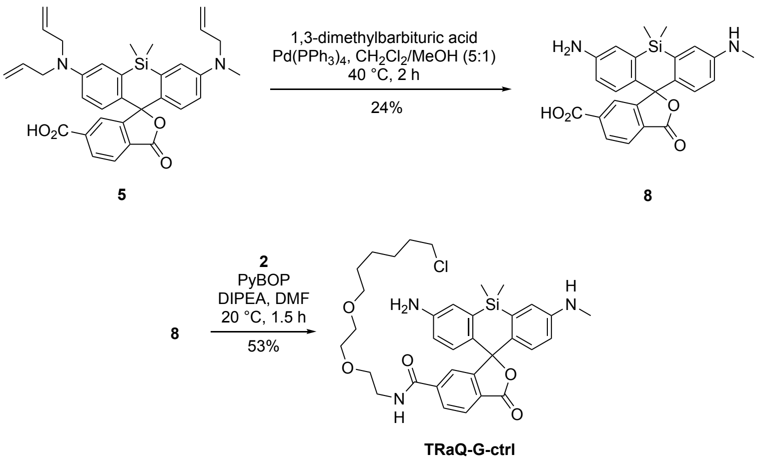

**Scheme S3**. Synthesis of the **TRaQ-G-ctrl** ligand.

3-Amino-5,5-dimethyl-7-(methylamino)-3'-oxo-3'$H$,5$H$-spiro[dibenzo[$b$,$e$]siline-10,1'-isobenzofuran]-6'-carboxylic acid (**8**)

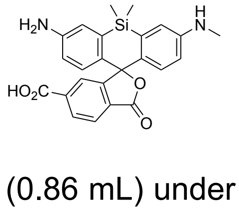

Compound **5**, 1,3-dimethylbarbituric acid (370 mg, 2.37 mmol, 20 equiv.) and tetrakis(triphenylphosphine)palladium(0) (68.5 mg, 59.2 µmol, 0.5 equiv.) were dissolved in a degassed mixture (5:1) of dichloromethane (0.42 mL) and methanol (0.86 mL) under an Ar atmosphere. The resulting mixture was stirred at 40 °C for 2 h. The reaction was diluted with $CH_2Cl_2$ and washed with saturated aqueous $Na_2CO_3$ solution. The aqueous phase was re-extracted with $CH_2Cl_2$ (2x). The combined organic phases were dried over $Na_2SO_4$ and the solvent was evaporated under reduced pressure. The crude product was purified by preparative HPLC (10 → 95% MeCN in dd$H_2O$ with 0.1% TFA) to yield a blue solid (9.3 mg, 21 mmol, 24%).
$^1$H NMR (800 MHz, MeOD) $\delta$ 8.30 (dd, $J$ = 8.1, 1.5 Hz, 1H), 8.24 (s, 1H), 7.86 (d, $J$ = 1.5 Hz, 1H), 7.26 (s, 1H), 7.14 (s, 1H), 6.89 (d, $J$ = 9.1 Hz, 1H), 6.69 (s, 2H), 6.61 (dd, $J$ = 9.2, 2.6 Hz, 1H), 2.98 (s, 3H), 0.64 (s, 3H), 0.56 (s, 3H).
$^{13}$C NMR (201 MHz, MeOD) $\delta$ 172.35, 168.02, 162.65, 150.50, 149.51, 136.53, 136.46, 136.42, 133.87, 133.41, 131.26, 131.21, 130.88, 129.78, 129.58,

126.30, 120.27, 120.16, 117.41, 117.27, 116.48, 115.22, 115.16, 100.02, 56.02, 54.20, 40.42, 38.99, -0.42, -1.69.
HRMS (ESI/QTOF) $[M+H]^+$ calculated for $C_{24}H_{23}N_2O_4Si^+$ 431.1422; found 431.1424.

3-Amino-*N*-(2-(2-((6-chlorohexyl)oxy)ethoxy)ethyl)-5,5-dimethyl-7-(methylamino)-3'-oxo-3'*H*,5*H*-spiro[dibenzo[*b,e*]siline-10,1'-isobenzofuran]-6'-carboxamide (**TraQ-G-ctrl**)

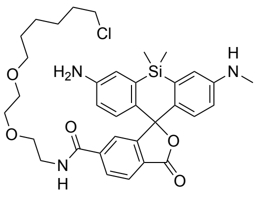

Chloroalkane **2**, compound **8** (7.0 mg, 16 µmol, 1 equiv.) and PyBOP (17 mg, 32 µmol, 2 equiv.) were added to a pre-dried flask under an $N_2$ atmosphere, dissolved in *N*,*N*-dimethylformamide (0.8 mL) and treated with DIPEA (27 µL, 0.16 mmol, 10 equiv.). The resulting solution was stirred at 20 °C for 2.5 h. All volatiles were removed, and the residue was purified by preparative HPLC (10 → 95% MeCN in $H_2O$ with 0.1% TFA) to yield a blue solid (5.7 mg, 8.6 mmol, 53%).
$^1$H NMR (600 MHz, MeOD) $\delta$ 8.77 (t, *J* = 5.2 Hz, 1H), 8.07 (s, 2H), 7.72 (d, *J* = 1.1 Hz, 1H), 7.10 (d, *J* = 2.5 Hz, 1H), 6.98 (d, *J* = 2.6 Hz, 1H), 6.69 (dd, *J* = 22.9, 8.8 Hz, 2H), 6.59 (dd, *J* = 8.7, 2.6 Hz, 1H), 6.51 (dd, *J* = 8.9, 2.7 Hz, 1H), 3.65 (t, *J* = 5.3 Hz, 2H), 3.63 – 3.60 (m, 2H), 3.59 – 3.54 (m, 4H), 3.51 (t, *J* = 6.7 Hz, 2H), 3.41 (t, *J* = 6.5 Hz, 2H), 2.83 (s, 3H), 1.73 – 1.66 (m, 2H), 1.49 (p, *J* = 6.7 Hz, 2H), 1.42 – 1.36 (m, 2H), 1.30 (p, *J* = 7.7 Hz, 2H), 0.63 (s, 3H), 0.54 (s, 3H).
$^{13}$C NMR (151 MHz, MeOD) $\delta$ 171.21, 168.68, 151.79, 149.84, 141.02, 140.23, 133.50, 133.24, 131.70, 130.83, 130.16, 129.25, 127.95, 127.60, 125.61, 121.58, 118.88, 117.41, 115.95, 114.31, 72.13, 71.19, 71.12, 70.32, 45.69, 41.13, 40.40, 33.70, 30.41, 30.29, 27.66, 26.40, -0.01, -1.47.
HRMS (ESI/QTOF) $[M+H]^+$ calculated for $[C_{34}H_{43}ClN_3O_5Si]^+$ 636.2655; found 636.2667.

### 5 NMR spectra

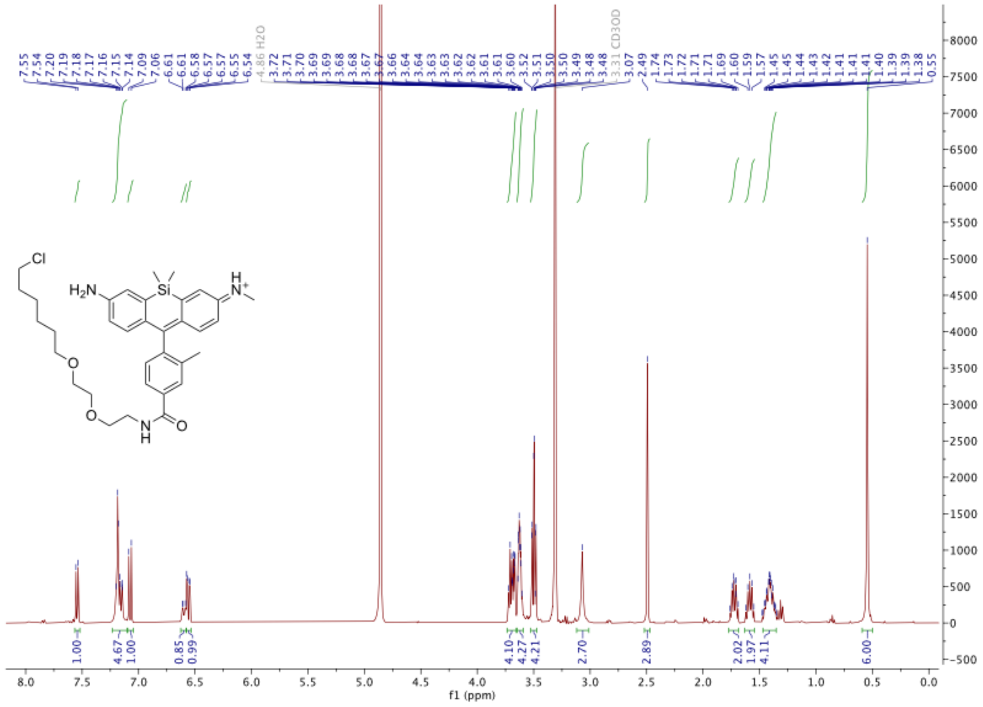

Figure S7. $^{1}$H-NMR of the **Me-TRaQ-G** ligand.

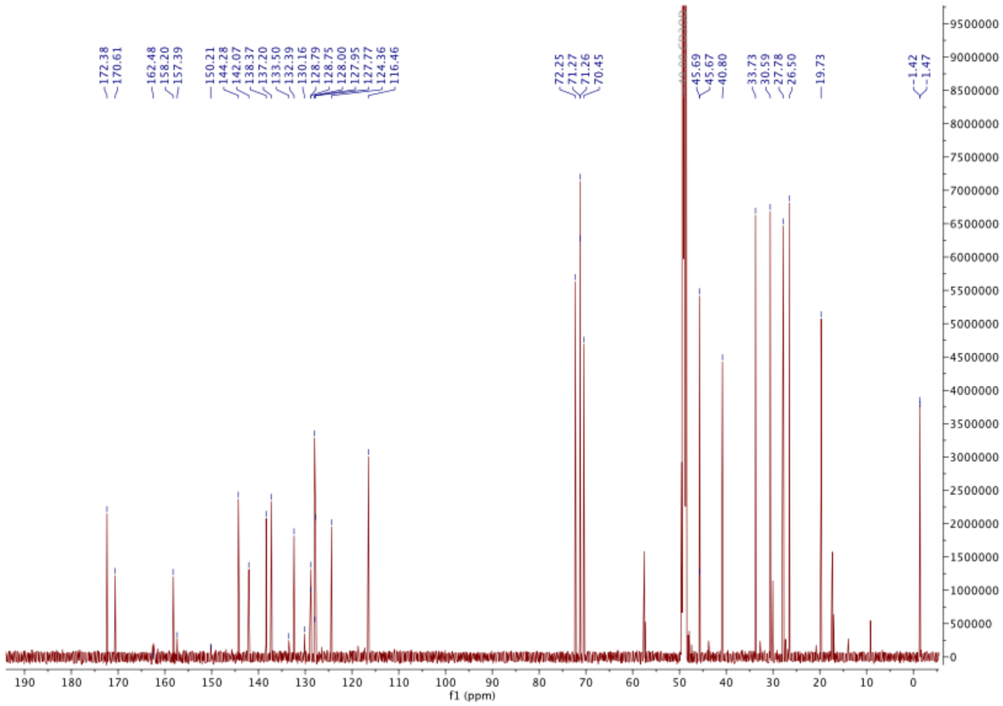

**Figure S8**. $^{13}$C-NMR of the **Me-TRaQ-G** ligand.

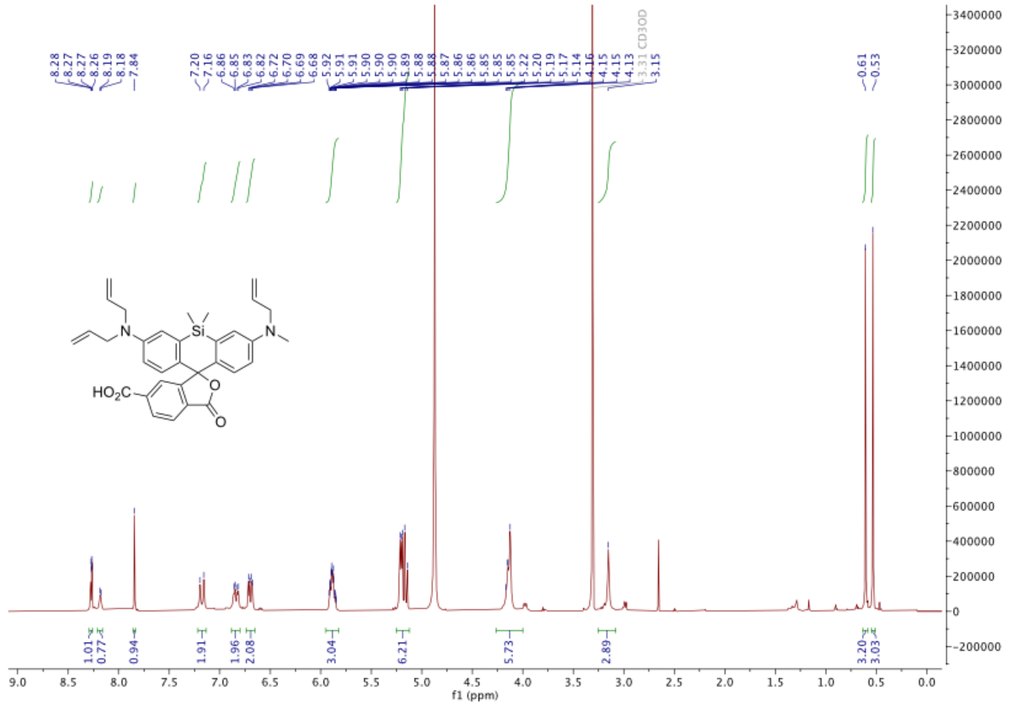

**Figure S9**. $^{1}$H-NMR of **5**.

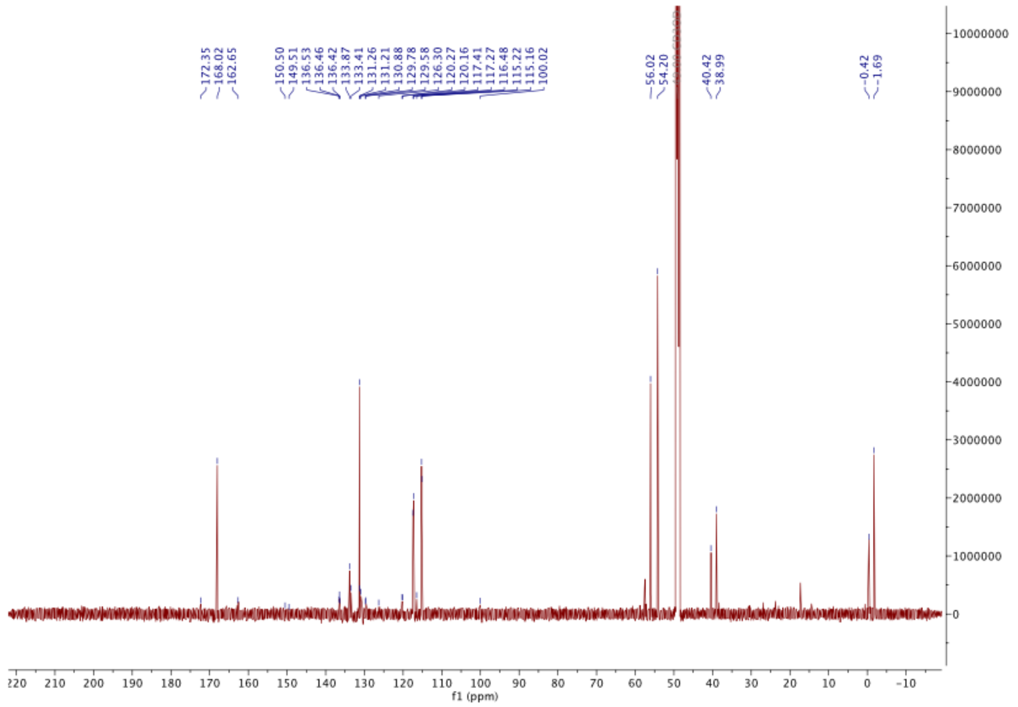

**Figure S10.** $^{13}$C-NMR of **5**.

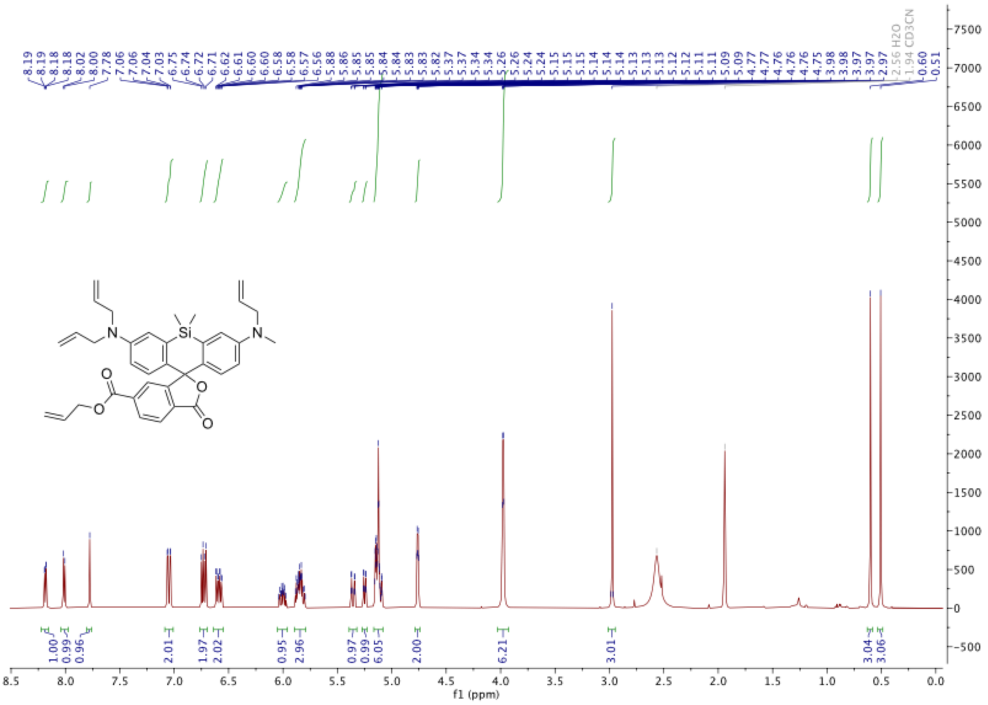

**Figure S11**. [1]H-NMR of **6**.

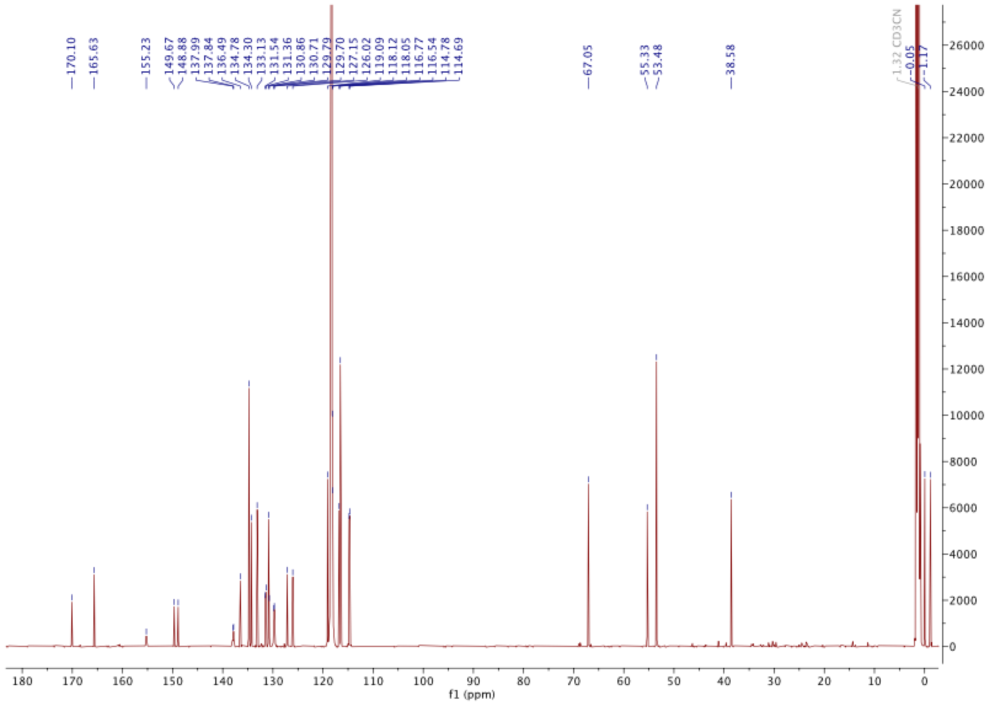

**Figure S12**. [13]C-NMR of **6**.

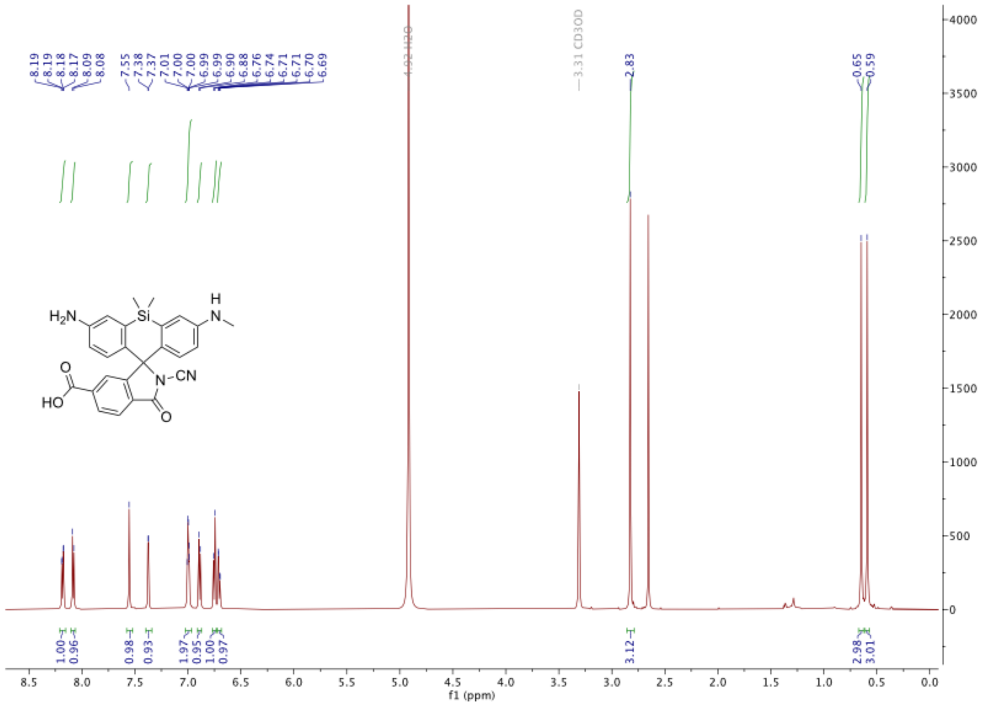

**Figure S13**. [1]H-NMR of **7**.

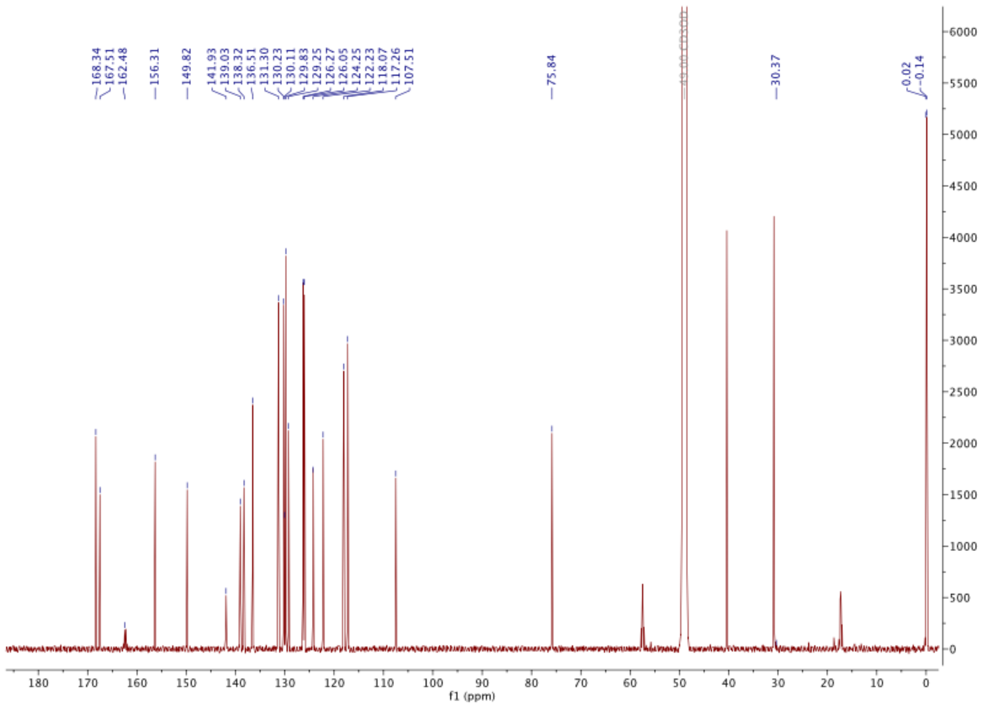

**Figure S14**. [13]C-NMR of **7**.

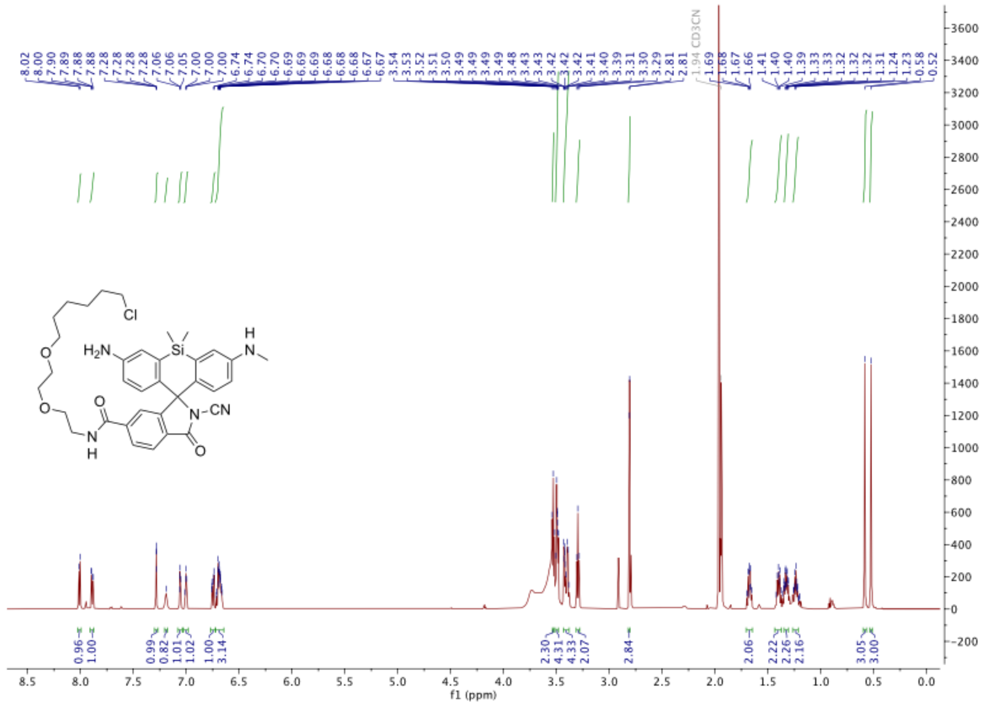

**Figure S15**. $^{1}$H-NMR of the **TRaQ-G** ligand.

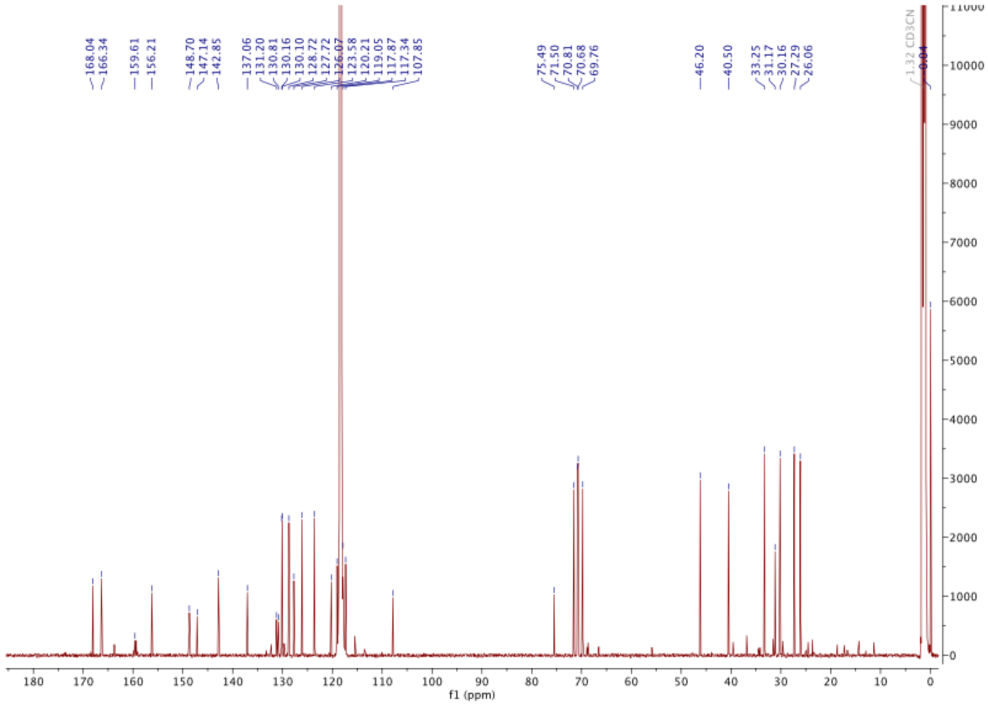

**Figure S16**. $^{13}$C-NMR of the **TRaQ-G** ligand.

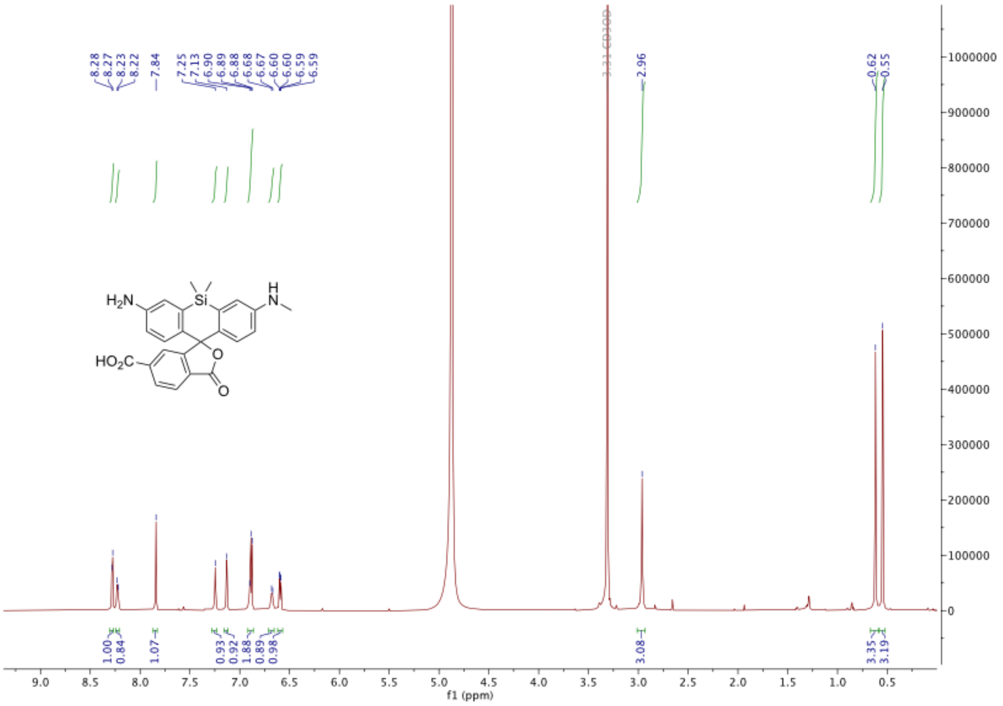

**Figure S17**. $^{1}$H-NMR of **8**.

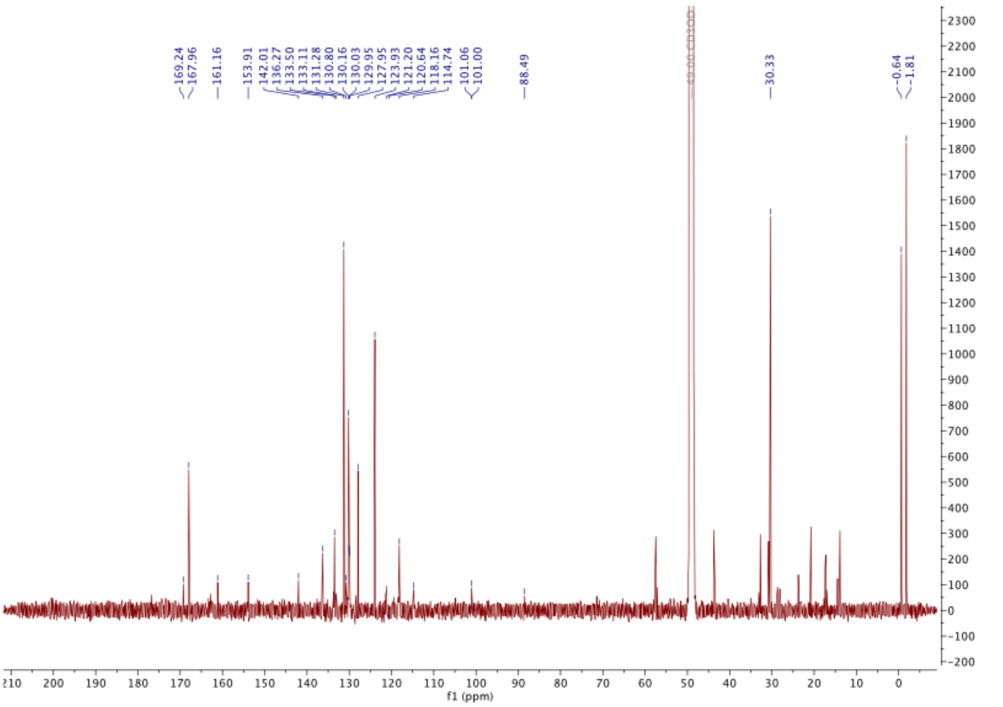

**Figure S18**. $^{13}$C-NMR of **8**.

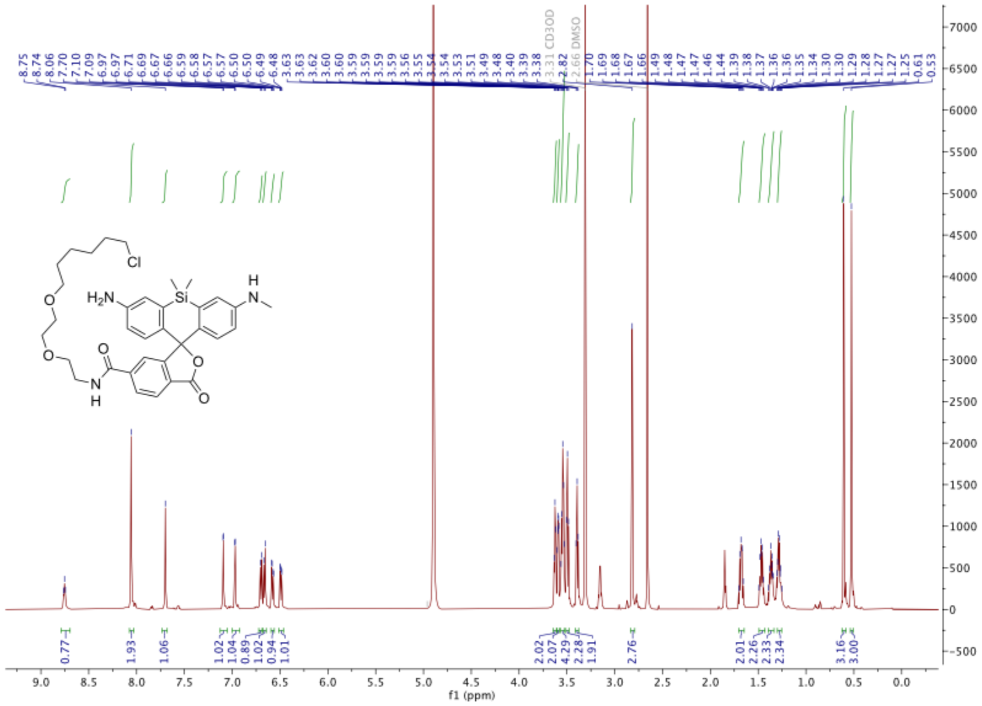

**Figure S19**. $^{1}$H-NMR of the **TRaQ-G-ctrl** ligand.

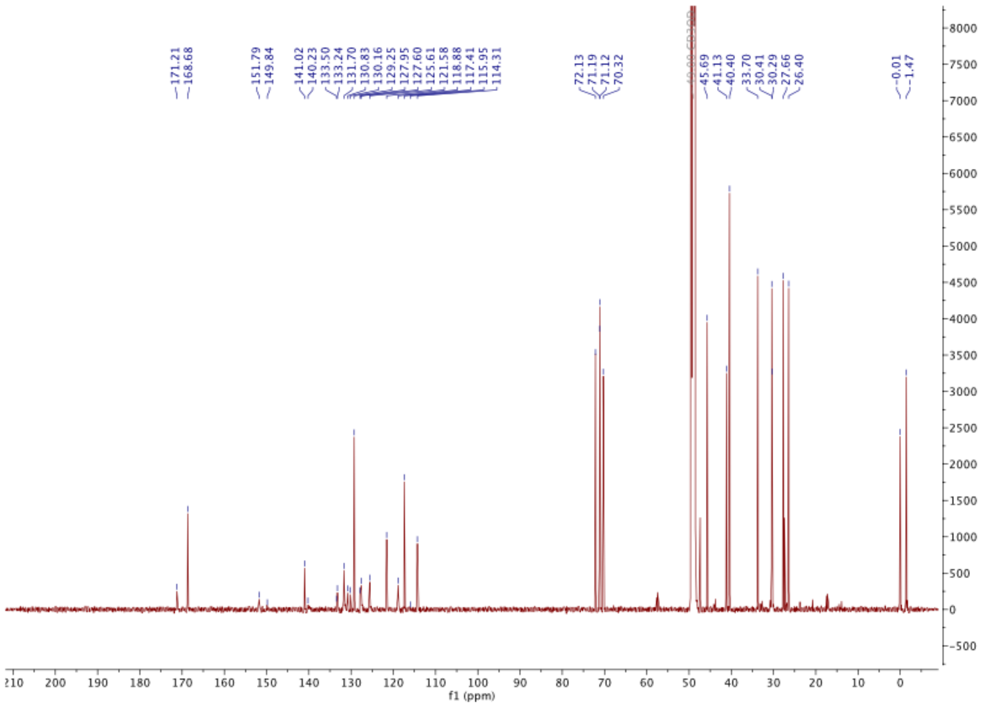

**Figure S20**. $^{13}$C-NMR of the **TRaQ-G-ctrl** ligand.
